# The Therapeutic Effects of Long-term Photobiomodulation on Aging in Mice

**DOI:** 10.1101/2023.08.05.552116

**Authors:** Ismayil Ahmet, Sunayana Begum Syed, Khalid Chakir, Christopher H Morrell, Praveen R Arany, Edward G Lakatta

## Abstract

**Background:** We have reported that photobiomodulation (PBM) therapy, a form of low dose Near Infrared Light (NIR) therapy, attenuates cardiovascular remodeling and extends the lifespan in a mouse model of accelerated cardiac aging. Here, we tested whether long-term PBM affects the aging process in normal male and female mice.

**Methods:** C57 mice, 18 months old, males (n=60) and females (n=60), were exposed to either NIR (850nm) at 25 mW/cm^2^ for 2 min on weekdays (MT and FT groups) or nothing (M and F groups) for 12 months. Mice were subjected to bimonthly echocardiography examination, Gait analysis and Frailty assessments. Randomly selected mice were sacrificed bimonthly for fresh tissue samples.

**Results:** Age-associated deterioration in left ventricle, left atrium, aorta, brain blood perfusion, frailty, body temperature and gait that were observed in M and F groups during the 12-month observation period were significantly attenuated by PBM therapy in MT and FT groups. The medium lifespan was extended by 0.6 and 1.0 month in MT and FT groups, compared to M and F groups, respectively. There was a significantly lower prevalence of dermatitis, stroke and heart failure in MT and FT groups compared to M and F groups.

**Conclusion:** Our data showed for the first time that PBM therapy by whole body exposure, even started at old age in normal animals, significantly attenuated the age-associated deterioration in heart, vessels, brain, gait and frailty; reduced the prevalence of stroke and heart failure; and improved health span.

## INTRODUCTION

The beneficial effects of sun light in human health are well known since ancient times. Accumulating evidence has shown the importance of sunlight in various diseases including cardiovascular health [1]. Recent advances in material physics helped to identify the wavelength of sun light (600-1000nm) that is most relevant to its beneficial effects and helped to birth of a light treatment, Photobiomodulation therapy (PBM) [2, 3]. PBM is a form of light therapy that utilizes non-ionizing forms of light sources, including Lasers, LEDs and broad-band light, in the visible and infrared spectrum [4, 5]. It is a non-thermal process involving endogenous chromophores eliciting photophysical and photochemical events at various biological scales.

Recently, we have reported that PBM attenuates cardiovascular remodeling and extends the lifespan in a mouse model of accelerated cardiac aging [6]. During the past decades, there has been a tremendous progress in understandings, application and mechanism of PBM. From isolated cells to whole animal studies, from preclinical to clinical studies, PBM has been shown to have remarkable beneficial therapeutic effects in wide variety of ailments [7–15]. In particular, the therapeutic effects of PBM treatments in age-associated neurocognitive and skeletomuscular diseases has been unequivocal such as Alzheimer’s disease, Parkinson’s disease, macular degeneration, chronic wounds, stroke and myocardial infarction among others [16–19]. Those beneficial effects of PBM have been broadly attributed to improvements in *in-situ* reprogramming of tissue-resident stem cells [20], in activation of cellular ATPase, in regulation of cytochrome c oxidase activity [21] and mitochondrial energy hemostasis [22], in activation of endogenous extracellular latent TGFβ signaling [23, 24] and NO signaling [25] via photoabsorption, non-visual photoreceptors and transponders such as TRPV1 and Opsin 2-5 [26–28]. Although key mechanistic pathways of PBM are still unknown, its safety, simplicity and effectiveness propelled its acceptance as an emerging new therapy for specific applications such as onco-therapy associated mucositis [29], wound healing, muscle pain, dentistry and others. Aging became one of the major social and medical issues that our societies are facing now.

Serving as a fertile ground for most of the major medical ailments like cancer, stroke, heart failure, diabetes, dementia and Alzheimer’s disease among elderly population, aging is and will be a huge burden to our society and health care system in foreseeable future. There is an urgent need to find a cure for aging. Because PBM alleviates most age-associated diseases, there is a real possibility that PBM may affect aging process itself. In this study, we focused on cardiovascular aging and investigated such possibility in C57 mice, a well characterized mouse strain that is widely used as a normal phenotype and serves as a common wild type background for most of the transgenic mouse strains.

In this proof of concept study, we initiated PBM treatment in C57 mice at 18 months of age to target the most dramatic phase of the aging that occurs during the second half of lifespan. We hypothesized that PBM treatment during the critical phase of cardiovascular aging may attenuate the age-associated cardiovascular remodeling, improve gait and frailty, and thereby improve health span and lifespan.

## METHODS

### Animal Studies

All animal studies were performed in accordance with the *Guide for the Care and Use of Laboratory Animals* published by the National Institutes of Health (NIH Publication no. 85-23, revised 1996). Experimental protocols were approved by the Animal Care and Use Committee of the National Institutes of Health (protocol #441LCS2022). C57 mice (n=120), males and females, 18 months old, were obtained from NIA Rodent Aging Colony and used in this study. Mice were housed in a climate-controlled room with 12-hour light cycle and free access to food and water. The study diagram was shown in Fig.1. Mice were randomly divided into 4 groups: (1) males without treatment (M group, n=30); (2) males with PBM treatment (MT group, n=30); (3) females without treatment (F group; n=30); and (4) females with PBM treatment (FT group, n=30). Mice were treated with PBM and observed for 12 months as described below. Mice were subjected to bimonthly echocardiography examination, Gait analysis and Frailty assessment. Mice were randomly selected for sacrifice bimonthly for fresh tissue samples (n=2 from each group for each time point). All survivals were sacrificed at the end of 12-month observational period.

**Figure 1:**
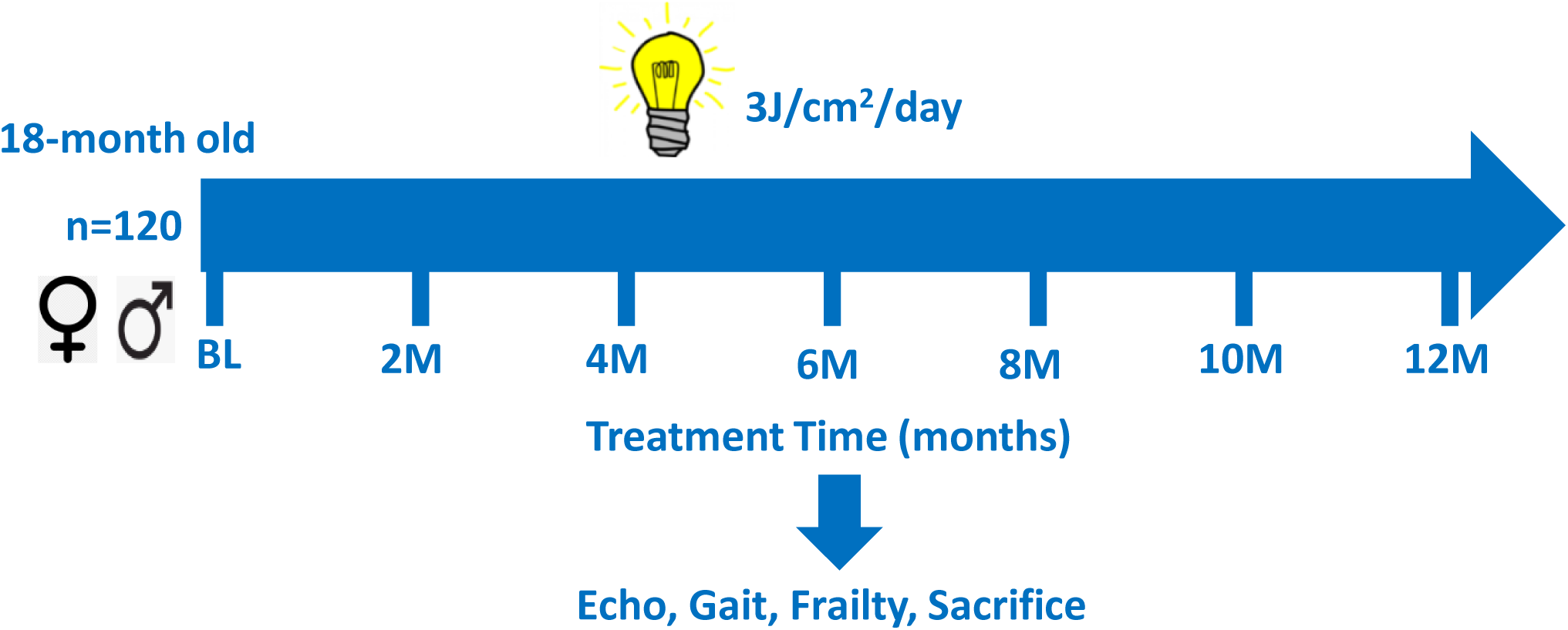
Study design. Light bulb indicates the presence of daily PBM treatment. C57 mice, male and female, n=120, 18 months old, were divided into 4 groups. Half of the mice were treated with daily dose of PBM for next 12 months. Every two months, mice were subjected to repeated tests and randomly selected for sacrifice.

### Photobiomodulation therapy

Mice in MT and FT groups received near-infrared LED light exposure at 25 mW/cm^2^ for 120s at 850 nm for 1 Einstein (4.5 p.J/cm^2^ photonic fluence or the conventional dose of 3 J/cm^2^) for 5 days per week for 12 months. Mice were placed under a LED light source (Bio900, Platinum LED Inc., Tampa, FL) in a fixed distance in their original cages without cage cover. Light irradiance was measured with an optical power meter (ThorLabs Inc., NJ) and adjusted accordingly. The mice were subjected to a whole-body treatment while they were awake and freely moving in their cage. As the PBM device was placed on the top of the cage, the dorsum (back) surface of the mice was directly exposed to the light. While shaving and restraining animals would have been ideal for PBM dosimetry, it was not considered appropriate from an animal use and care perspective due to the repeated, extensive, long-term treatment regimen. To ensure suitable dosimetry, average irradiance in ambulatory animals at a median anatomical plane was calculated. Mice were treated at a same time each day between 10 and 11 am. Mice in M and F groups went through same procedure without light exposure.

### Cardiovascular assessment

All mice in the study underwent bimonthly echocardiography (Echo) examination (40-MHz transducer; Visual Sonics 3100; Fuji Film Inc, Seattle, WA) under light anesthesia with isoflurane (2% in oxygen) via nosecone and were kept at 37°C using a heating pad. Mice were placed in the supine position; skin hair in the chest area was shaved.

Body weight (BW) was measured. Standard ECG electrodes were placed on the limbs and Lead II ECG was recorded with echo images. Heart rate (HR) was calculated from 5 consecutive RR intervals of ECG recordings. Each Echo examination was completed within 10 min. Parasternal long-axis views of the left ventricle (LV) were obtained and recorded to ensure that the mitral and aortic valves and the LV apex were visualized. M-mode tracing of Left atrium and basal aorta were recorded at aortic valve level. Left atrium end-systolic dimension (LAD) and Aortic end-systolic dimension (AoD) were measured and LADc was calculated as LAD/AoD. Parasternal short-axis views of the LV were recorded at the mid-papillary muscle level. From the parasternal short-axis view of the LV, M-mode tracings of the LV were obtained. Thicknesses of LV posterior Wall (PW) and intraventricular septum (IVS) were measured from M-mode tracing of LV. Endocardial area tracings, using the leading-edge method, were performed in the 2D mode (short-axis and long-axis views) from digital images captured on cine loop to calculate the end-diastolic and end-systolic LV areas. LV End-diastolic volume (EDV) and end-systolic volume (ESV) were calculated by a Hemisphere Cylinder Model method. EF was derived as EF = 100 * (EDV-ESV) / EDV. Stroke volume (SV) was the net difference between EDV and ESV. Cardiac Output (CO) calculated as CO = SV * HR. Cardiac Index (CI) calculated as CI = CO/BW. LV mass (LVM) was calculated from EDV, IVS and PWth. Four chamber view of heart was obtained from the apex. Mitral valve blood flow velocity was recorded at the tip of the mitral valves using pulsed Doppler. E and A wave velocities and E/A ratio were measured off-line. Mitral valve Tissue Doppler velocity was recoded at the fibrotic ring of mitral valve at IVS. e’ and a’ wave velocities and e’/a’ ratio were measured off-line. A 2D image of Thoracic aorta at long axis was recorded right above diaphragm. Thoracic aortic lumen dimensions at maximum and minimum (TAmax; TAmin) were measured. A fractional changes of lumen dimension during a cardiac cycle was calculated as TAFC = (TAmax – TAmin) * 100 / TAmin. Pulse wave velocity (PWV) was measured at lower part of abdominal aorta. Doppler flow velocity was acquired at 2 points of abdominal aorta: just above to the iliac bifurcation and 6 ∼ 10mm proximal to the iliac bifurcation. The distance between two sampling points was measured from 2D images using Image J software (NIH). The time between R wave of ECG and starting point of Doppler flow velocity wave was measured off-line for both sampling points. Their net difference was used as the wave traveling time between two points. PWV was calculated as PWV = distance (mm) / traveling time (s). A 2D image of Carotid artery at long axis and a Doppler flow velocity were recorded right before carotid bifurcation. Carotid artery lumen dimensions at maximum and minimum (CAmax; CAmin) were measured; Blood flow (CAFlow) was calculated from carotid artery lumen dimensions and Doppler flow velocity VTI tracings. A fractional changes of lumen dimension during a cardiac cycle was calculated as CAFC = (CAmax – CAmin) * 100 / CAmin. All measurements were made by a single observer who was blinded to the identity of the tracings. All measurements were an average of five consecutive cardiac cycles that covering at least one respiration cycle. The reproducibility of measurements was assessed by one repeated measurement that week apart in randomly selected images, and the repeated-measure variability was less than 5%.

### Gait analysis

All mice in the study underwent bimonthly Gait examination (DigiGait System, Mouse Specifics Inc. Framingham, MA). DigiGait ™ was used to analyze gait parameters per the manufacturer’s protocol. Recording of gait and analysis of gait were described in details previously [30]. Briefly, at each time point, unconstrained mouse was put on the treadmill belt for 1 min for acclamation and forced to run at a belt speed of 5cm/s for 2 min, 15cm/s for 5 min, and then 18 cm/s while recording. When mouse has difficulties running or refused to run, they would be encouraged through reversing the direction of the belt, gently prodding with the bumpers, or removal from the chamber to reattempt after a few minutes of rest. If mouse still can’t cooperate, the maximum belt speed it could run was used for recording. After obtaining a 4∼6 sec of video recording of steady run, mouse was returned to original cages. Gait parameters were analyzed off-line with the DigiGait ™ software (Mouse Specifics, Inc.).

Fifty-four parameters were evaluated for all four limbs through software automation and reported as an average of four limbs. Temporal, spatial, postural, intra-individual variability and nerve injury parameters were reported for each group. Temporal parameters are based on time and indicate fluidity of a leg movement. Gait Speed is a maximum voluntary walking speed, a parameter of limb skeletal muscle functional reserve. Spatial parameters are based on the paw area and indicate the balance and coordination among 4 limbs. Gait symmetry is an important spatial parameter. In a healthy subject walking fluidly, the gait symmetry will be in unity and close to 1. Postural parameters are based on the style of paw placement and indicate abnormalities of each limb. Animal Width is the average distance between right and left paws that reflects the walking posture. Ataxia Coefficient reflects a balance among four limbs during the walking and is an important parameter of brain-muscle coordination. Intra-individual variability parameters are based on the variation of each parameter from multiple work cycles and indicate functional stability of neuro-muscular coordination. Swing Duration CV is an important intra-individual variability parameter, reflects the dispersion about the average value.

Swing Duration CV indicates how regular the consecutive walking cycles is. The Sciatic Functional Index (SFI), the Tibial Functional Index (TFI) and the Peroneal Functional Index (PFI) of hind limbs indicate a presence of peripheral neuropathic pain, injury or dysfunction involving sciatic nerve, tibial nerve or peroneal nerve, respectively. All assessments were made by an investigator blinded to group identities.

### Frailty assessment

All mice in the study underwent bimonthly frailty assessment. The clinical frailty index was conducted as previously described [31]. In brief, 29 deficits were examined and graded with increasing severity, with each criterion scored 0, 0.5 or 1. Bodyweight and body temperature were analyzed separately and not included in the frailty score. The frailty index score was derived from the sum of deficits, divided by the total possible [32] deficits, resulting in a total mouse clinical frailty index score between 0 and 1. Mice were considered frail when above the published cut-off of 0.18 for the clinical frailty index [31,33].

### Animal sacrifice, tissue collection and autopsy

Randomly selected 2 mice from each group were sacrificed at every 2 months for fresh tissue samples. All surviving animals in the study were sacrificed at the end of 12-month observation period. A through autopsy was conducted on all subjects sacrificed and deceased by natural cause. Presence of dermatitis and stroke were determined by facility vet. Presence of heart failure was determined if EF<30% prior to death. All abnormal growth masses found during autopsy were classified as tumors and their samples were collected for further analysis. Organs were weighed and tissue samples from blood serum, skin, brain, lung, heart, aortas, liver, kidney, leg muscle were collected and stored at −80^0^C.

### Statistical Analyses

All data are expressed as mean ± SEM. Statistical significance was assumed at p<0.05. A Kaplan-Meier survival curve was computed for each group and a log-rank test applied to determine if the survival curves differ among groups. For data from repeated measurements, we applied a linear mixed-effects model to perform a 2-way mixed-ANOVA analysis on two main effects: treatment and time. It was done separately for males and females. Backward elimination of non-significant terms, starting with the highest order terms, yields the final model for each of the parameters. When any 2-way interaction (treatment x time) is significant, a post-hoc comparison with a FDR correction for multiple testing was used to determine at what time point it is differ. For data from single measurements, ANOVA with a Banforroni post-hoc test were applied.

## RESULTS

There were no exclusions in this study. All 120 mice were accounted for data analysis.

### Cardiac Chamber Remodeling

EDV and ESV are the parameters of LV chamber size. SV is the net difference between EDV and ESV. The changes of EDV, ESV and SV during the 12-month observation period were shown in Fig.2. In males, EDV and ESV were gradually increased by age, while SV was gradually reduced by age, indicate an on-set of age-associated LV chamber dilation and also indicate that LV functional deterioration (increases in ESV) over takes LV structural remodeling (increases in EDV). PBM significantly altered the trajectories of all 3 parameters; it significantly reduced EDV and ESV and increased SV from their pre-treatment values, indicates that PBM reversed both LV functional deterioration and LV structural remodeling. In females, EDV, ESV and SV were gradually increased by age, indicate an on-set of age-associated LV chamber dilation and also indicate that LV structural remodeling (increases in EDV) over takes LV functional deterioration (increases in ESV). PBM significantly altered the trajectories of EDV and ESV, but not of SV; it significantly slowed down the increases in EDV and ESV at about a same rate, indicates that PBM attenuated LV functional deterioration and LV structural remodeling.

**Figure 2:**
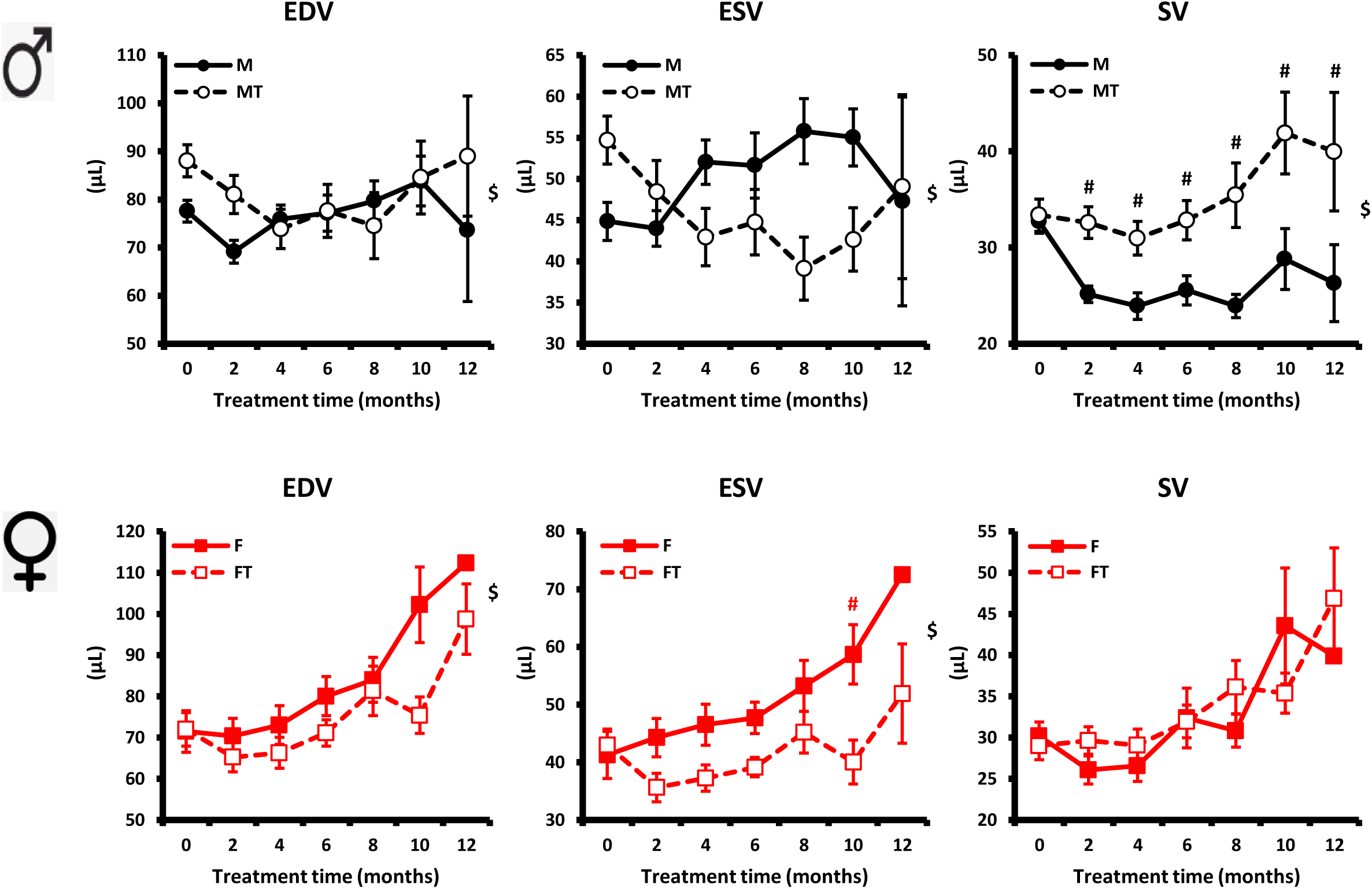
Cardiac parameters. EDV = LV end-diastolic volume; ESV = LV end-systolic volume; EF = LV ejection fraction. SV = LV stroke volume. CO = Cardiac output. CI = Cardiac Index. Mean ± SEM; $ p<0.05 for Group-time interaction; post-hoc test: # (red) p<0.05 F vs. FT; # (black) p<0.05 M vs. MT.

### Cardiac Function

EF, CO and CI are the parameters of LV function. The changes of EF, CO and CI during the 12-month observation period were shown in Fig.3. In males, EF, CO and CI were gradually reduced by age; indicate an on-set of age-associated cardiac functional deterioration and development of a systolic heart failure. PBM significantly altered the trajectories of all 3 parameters; it significantly increased EF, CO and CI from their pre-treatment values, indicates that PBM reversed such LV functional deterioration and prevented heart failure. In females, EF was gradually reduced by age, while CO and CI were gradually increased by age, indicates an on-set of age-associated cardiac functional deterioration, but without development of a systolic heart failure. Such increases in CO and CI were largely attributed to the increases in SV and HR (see below). PBM significantly altered the trajectory of EF, but not of CO and CI; it significantly increased EF from its pre-treatment value, indicating PBM attenuated such LV functional deterioration.

**Figure 3:**
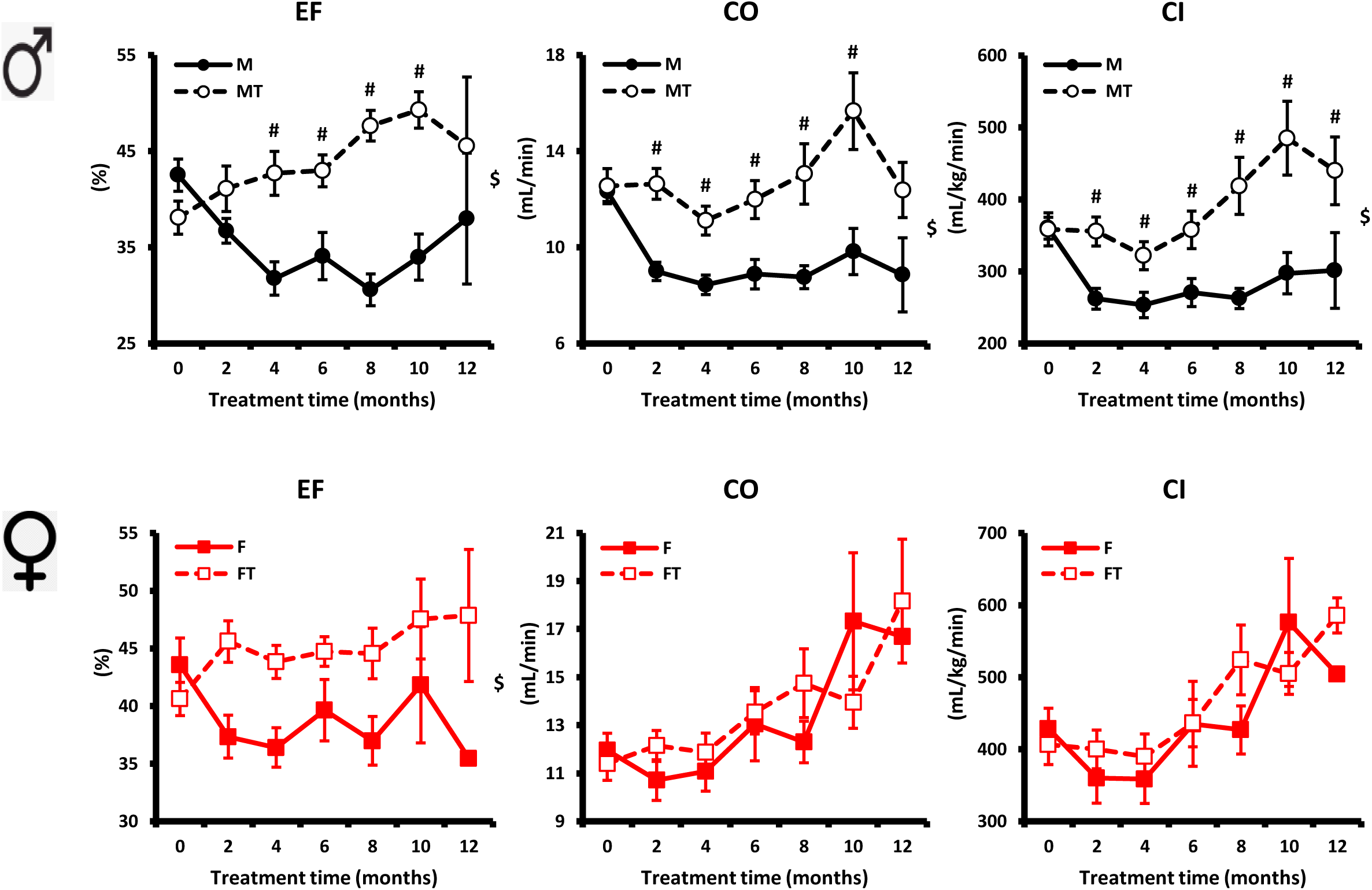
Cardiac parameters. EF = LV ejection fraction; CO = Cardiac output; CI = Cardiac Index. Mean ± SEM; $ p<0.05 for Group-time interaction; post-hoc test: # (red) p<0.05 F vs. FT; # (black) p<0.05 M vs. MT.

### Cardiac Hypertrophy

LVM is parameter of cardiac hypertrophy. It is calculated from EDV, IVS and PW. The changes of LVM and PW during the 12-month observation period were shown in Fig.4. In males, LVM and PW were gradually increased by age, indicates an on-set of age-associated cardiac hypertrophy. Such increase in LVM was mainly attributed to the increases in PW. PBM significantly altered the trajectory of LVM and attenuated the increases in LVM, indicates that PBM prevented cardiac hypertrophy. In females, LVM was gradually increased by age, indicates an on-set of age-associated cardiac hypertrophy. Such increase in LVM was mainly attributed to the increases in EDV. PBM significantly altered the trajectories of both PW and LVM and attenuated the increases in LVM and PW, indicates that PBM prevented cardiac hypertrophy.

**Figure 4:**
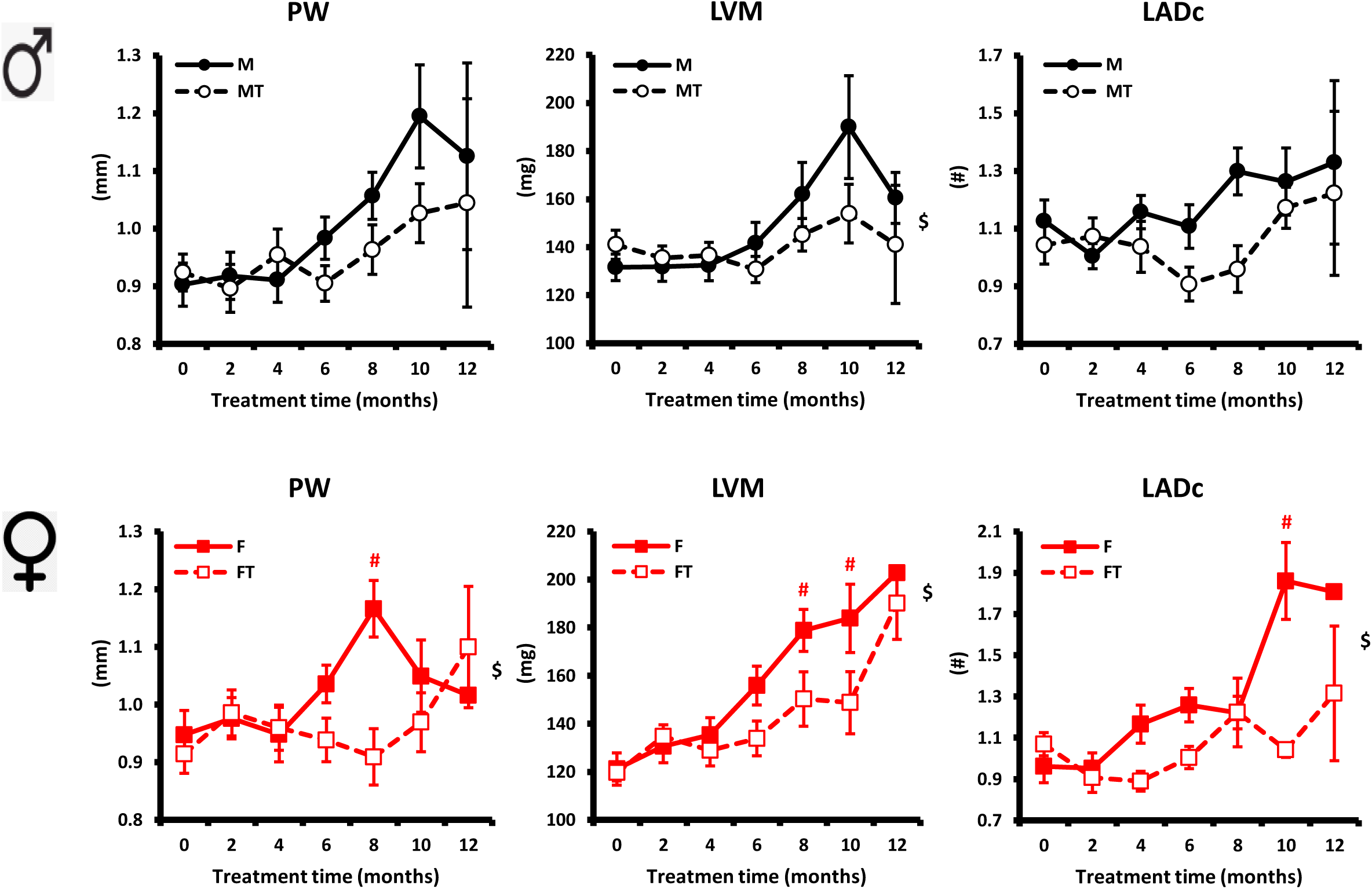
Cardiac parameters. PW = LV posterior wall thickness; LVM = LV mass; LADc = left atrial dimension adjusted to aortic lumen dimension. Mean ± SEM; $ p<0.05 for Group-time interaction; post-hoc test: # (red) p<0.05 F vs. FT; # (black) p<0.05 M vs. MT.

### Cardiac Diastolic Function

LADc, Mitral E/A ratio and Tissue Doppler e’/a’ ratio are the parameters of LV diastolic function. The changes of LADc during the 12-month observation period were shown in Fig.4. In both males and females, LADc was gradually increased by age, indicates an on-set of age-associated LV diastolic dysfunction. PBM significantly altered the trajectory of LADc in females; it significantly reduced LADc from its pre-treatment value in both male and females, indicates that PBM prevented cardiac diastolic dysfunction. The changes of Mitral E/A ratio and Tissue Doppler e’/a’ ratio were shown in Supplement Data (Supp. Fig.1). Both parameters were not significantly different between two groups in males and in females during the 12-month observation period.

### Aortic Wall Stiffness

PWV, TAFC and CAFC are the parameters of aortic wall stiffness. The changes of PWV, TAFC and CAFC during the 12-month observation period were shown in Fig.5. In males, PWV was gradually increased, and TAFC and CAFC were gradually reduced by age, indicates an on-set of age-associated aortic wall remodeling. PBM significantly altered the trajectories of all 3 parameters; it significantly attenuated increases in PWV and reductions in TAFC and CAFC, and also increased TAFC from its pre-treatment value, indicates that PBM prevented the aortic wall remodeling. In females, PWV was gradually increased, and TAFC and CAFC were gradually reduced by age, indicates an on-set of age-associated aortic wall remodeling. PBM significantly altered the trajectories of PWV; it significantly attenuated increases in PWV, indicates that PBM prevented the aortic wall remodeling.

**Figure 5:**
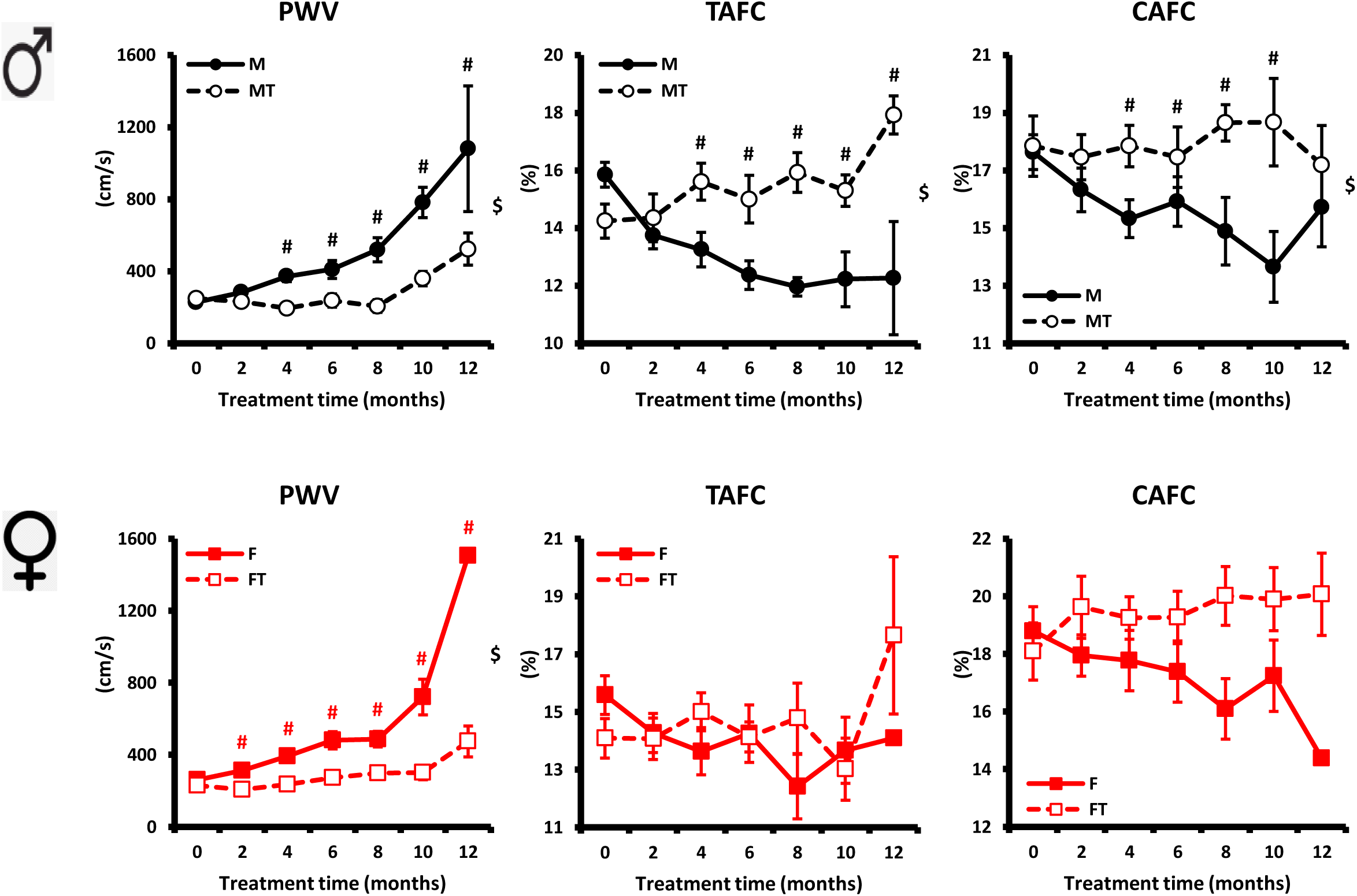
Vascular parameters. PWV = Aortic pulse wave velocity; TAFC = coronary artery lumen fractional changes; CAFC = coronary artery lumen fractional changes. Mean ± SEM; $ p<0.05 for Group-time interaction; post-hoc test: # (red) p<0.05 F vs. FT; # (black) p<0.05 M vs. MT.

### Carotid Blood Flow

CAFlow is an indirect indicator of cerebral blood perfusion level. CF/CO shows the portion (or percent) of total cardiac output goes to the brain. The changes of CAFlow and CF/CO during the 12-month observation period were shown in Fig.6. In males, CAFlow was gradually increased by age, indicates an on-set of age-associated increase in cerebral perfusion. CF/CO was also gradually increased by age while CO was reduced. It shows how brain is prioritized over rest of the body. PBM didn’t affect CAFlow. It significantly attenuated the increase in CF/CO, largely due to the preserved CO. In females, CAFlow was gradually increased by age, indicates an on-set of age-associated increase in cerebral perfusion. CF/CO was also gradually increased by age while CO was also increased. It shows the increase in cerebral perfusion by age outpaced the increases in CO. PBM attenuated the increase in CAFlow. It significantly reduced CF/CO from its pre-treatment value, largely due to the reduction in CAFlow.

**Figure 6:**
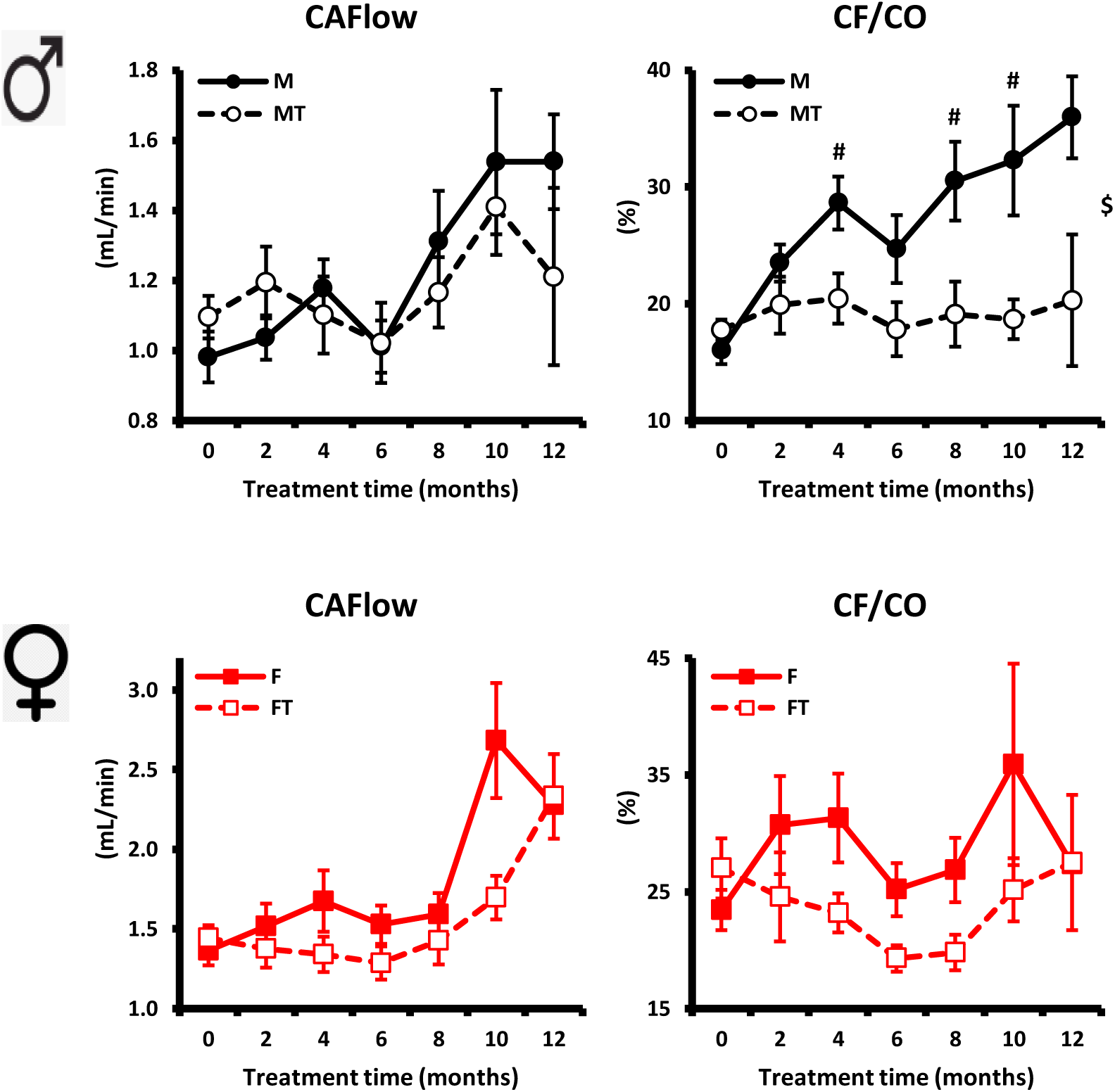
Brain blood perfusion parameters. CA Flow = right coronary artery blood flow; CF/CO = percent CA Flow to CO ratio. Mean ± SEM; $ p<0.05 for Group-time interaction; post-hoc test: # (red) p<0.05 F vs. FT; # (black) p<0.05 M vs. MT.

### Gait

The changes of the representative gait parameters during the 12-month observation period were shown in Fig.7. Rest of the parameters was shown in Supplement Data (Supp. Fig.2). In males, Gait Speed was gradually reduced by age; Swing Duration CV was increased by age. It indicates an on-set of age-associated deterioration of limb skeletal muscle reserve and walking rhythm and posture. Ataxia Coefficient did not change by age. PBM significantly altered the trajectories of Gait Speed and Swing Duration CV; indicates that PBM prevented skeletal muscle dysfunction and preserved youthful walking rhythm and posture. In females, Gait Speed was gradually reduced by age; Swing Duration CV was increased by age. It indicates an on-set of age-associated deterioration of limb skeletal muscle reserve and walking rhythm and posture. Ataxia Coefficient did not change by age. PBM significantly altered the trajectories of Gait Speed and Ataxia Coefficient; indicates that PBM prevented skeletal muscle dysfunction and preserved youthful walking rhythm, posture and balance.

**Figure 7:**
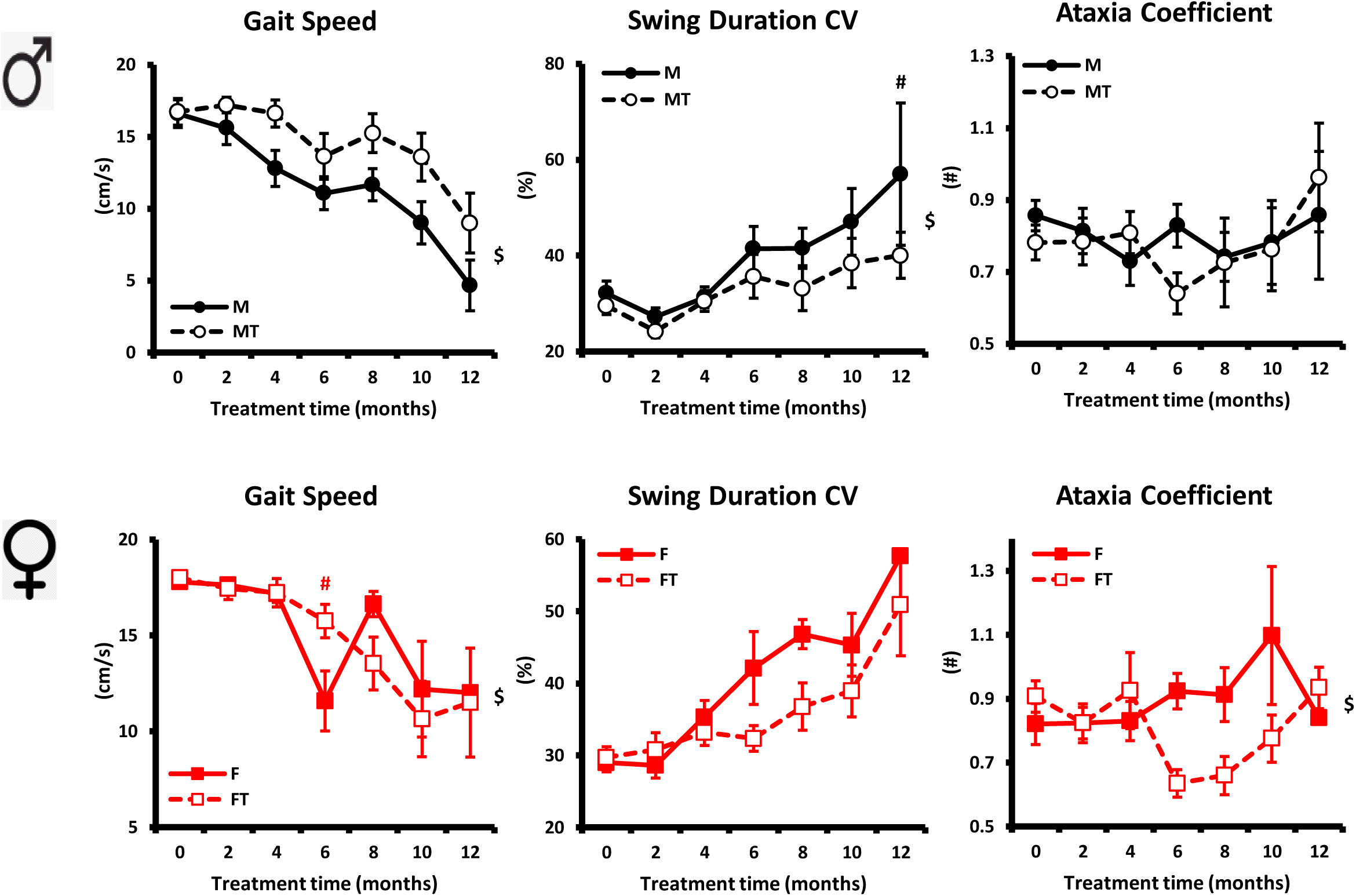
Gait parameters: Gait speed, Swing Duration CV and Ataxia coefficient. Mean ± SEM; $ p<0.05 for Group-time interaction; post-hoc test: # (red) p<0.05 F vs. FT; # (black) p<0.05 M vs. MT.

### Gait Peripheral Neuropathic Pain/injury/dysfunction Index

The SFI, TFI and PFI of hind limbs indicate a presence of peripheral neuropathic pain, injury or dysfunction involving sciatic nerve, tibial nerve or peroneal nerve, respectively. The changes of SFI, TFI and PRI during the 12-month observation period were shown in Fig.8. In males, all 3 parameters were gradually increased by age between 24 and 30 months of age; indicates an on-set of age-associated peripheral neuropathic pain/injury/dysfunction in advanced age. PBM significantly altered the trajectory of all 3 parameters; indicates that PBM prevented peripheral neuropathic pain/injury/dysfunction. In females, all 3 parameters did not changed by age or affected by PBM.

**Figure 8:**
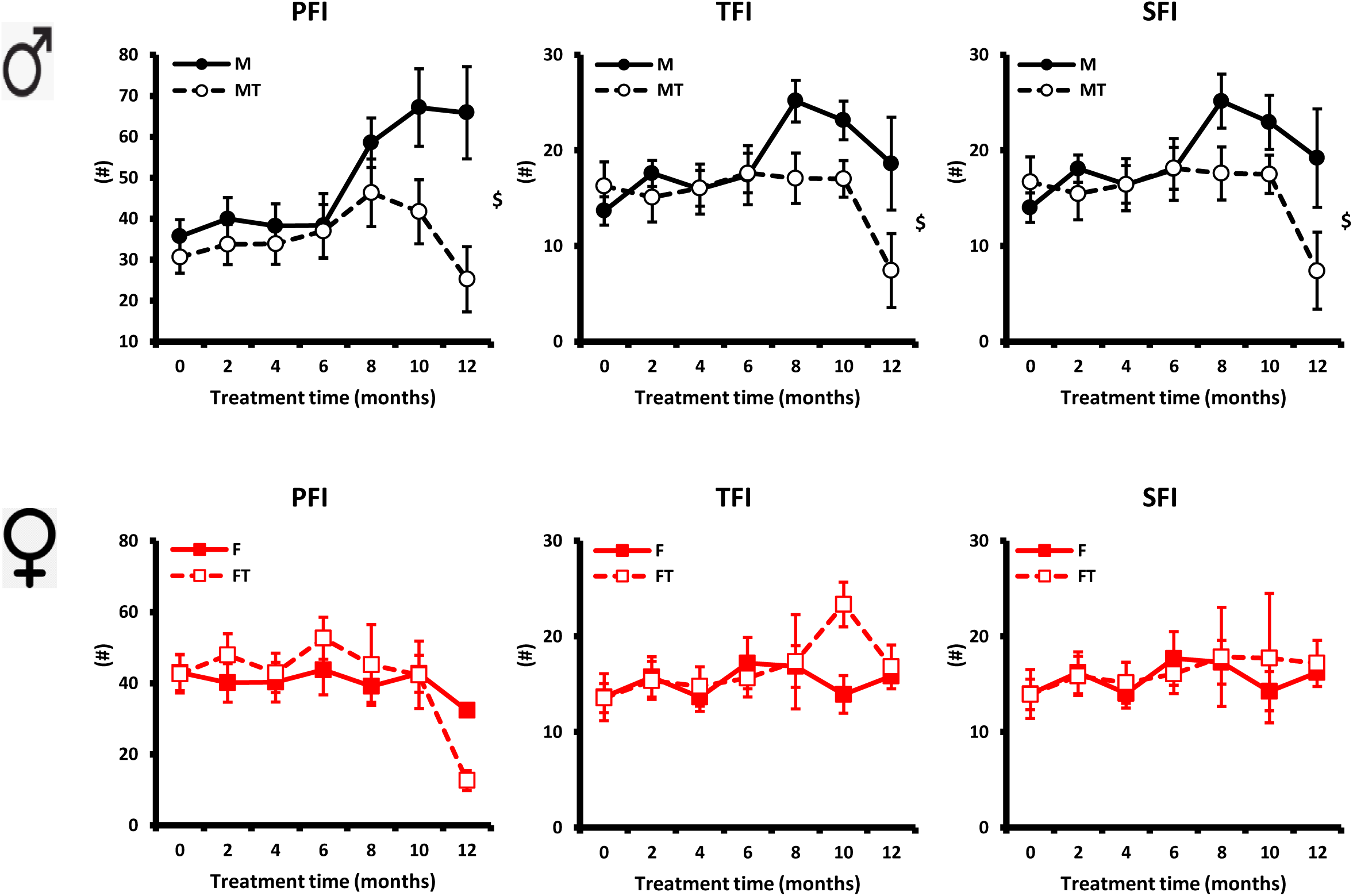
Gait Peripheral Neuropathic Pain/injury/dysfunction Indices. SFI = the sciatic functional index; TFI = the tibial functional index; PFI = the peroneal functional index. Mean ± SEM; $ p<0.05 for Group-time interaction; post-hoc test: # (red) p<0.05 F vs. FT; # (black) p<0.05 M vs. MT.

### Frailty

The changes of Frailty Score and Body Temperature during the 12-month observation period were shown in Fig.9. In males, Frailty Score was gradually increased by age; Body Temperature was gradually reduced by age, indicates an on-set of age-associated deterioration of health and thermoregulation. PBM significantly altered the trajectory of Frailty Score; and slightly increased Body Temperature; indicates that PBM preserved youthful health and improved thermoregulation. In females, Frailty Score was gradually increased by age; indicates an on-set of age-associated deterioration of health. Body Temperature did not change by age. PBM significantly altered the trajectories of Frailty Score and Body Temperature; indicates that PBM preserved youthful health and improved thermoregulation.

**Figure 9:**
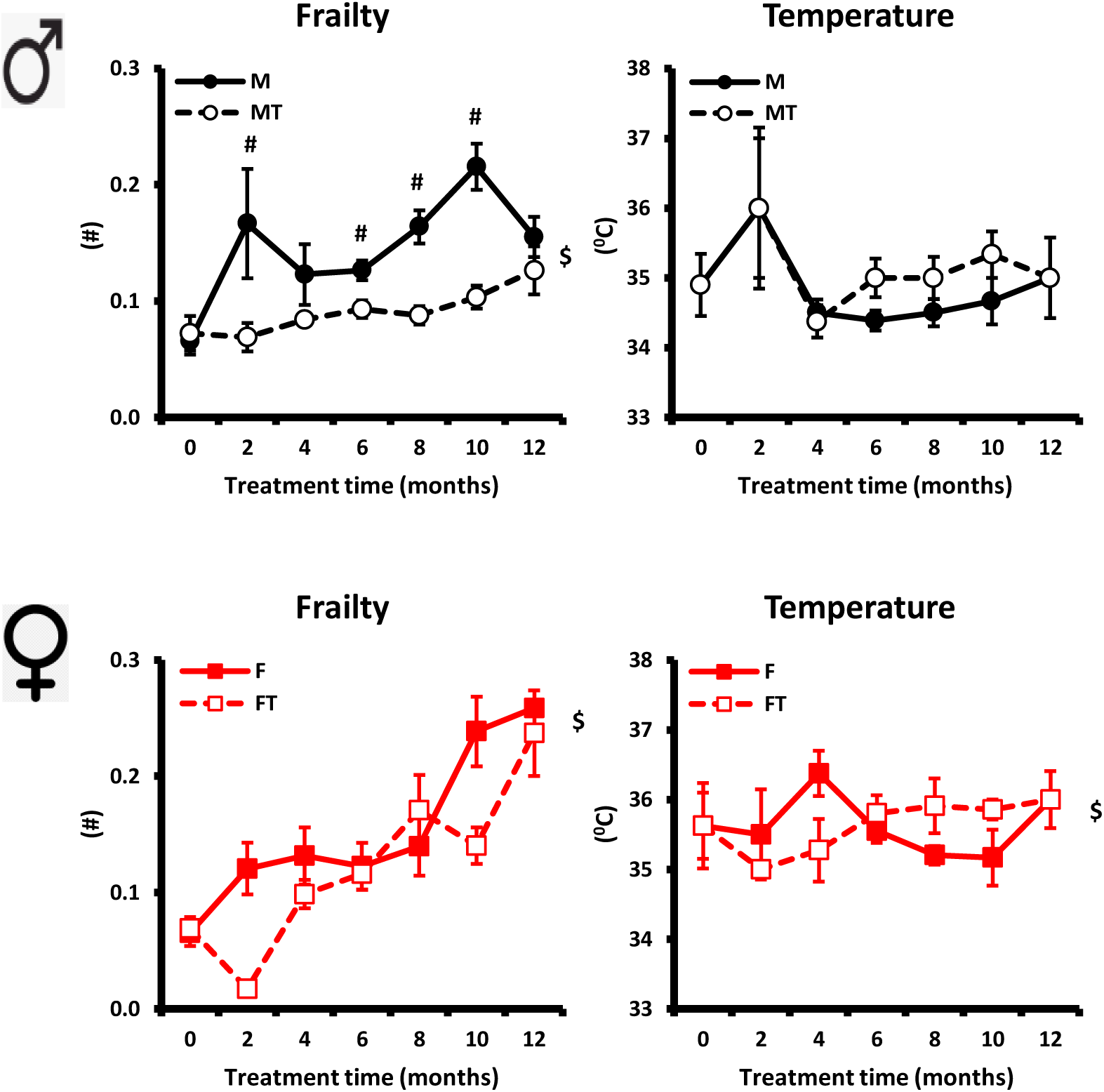
Frailty and Body surface temperature. Mean ± SEM; $ p<0.05 for Group-time interaction; post-hoc test: # (red) p<0.05 F vs. FT; # (black) p<0.05 M vs. MT.

### Body Weight and Heart Rate

The changes of BW and HR during the 12-month observation period were shown in Fig.10. In males, both BW and HR were gradually reduced by age, indicates an on-set of age-associated loss of body mass and deterioration in heart’s pacemakers. PBM did not affect BW; PBM slightly increased the HR, but it did not reach statistical significance. In females, both BW and HR showed a similar biphasic changes by age. PBM did not affect either BW or HR.

**Figure 10:**
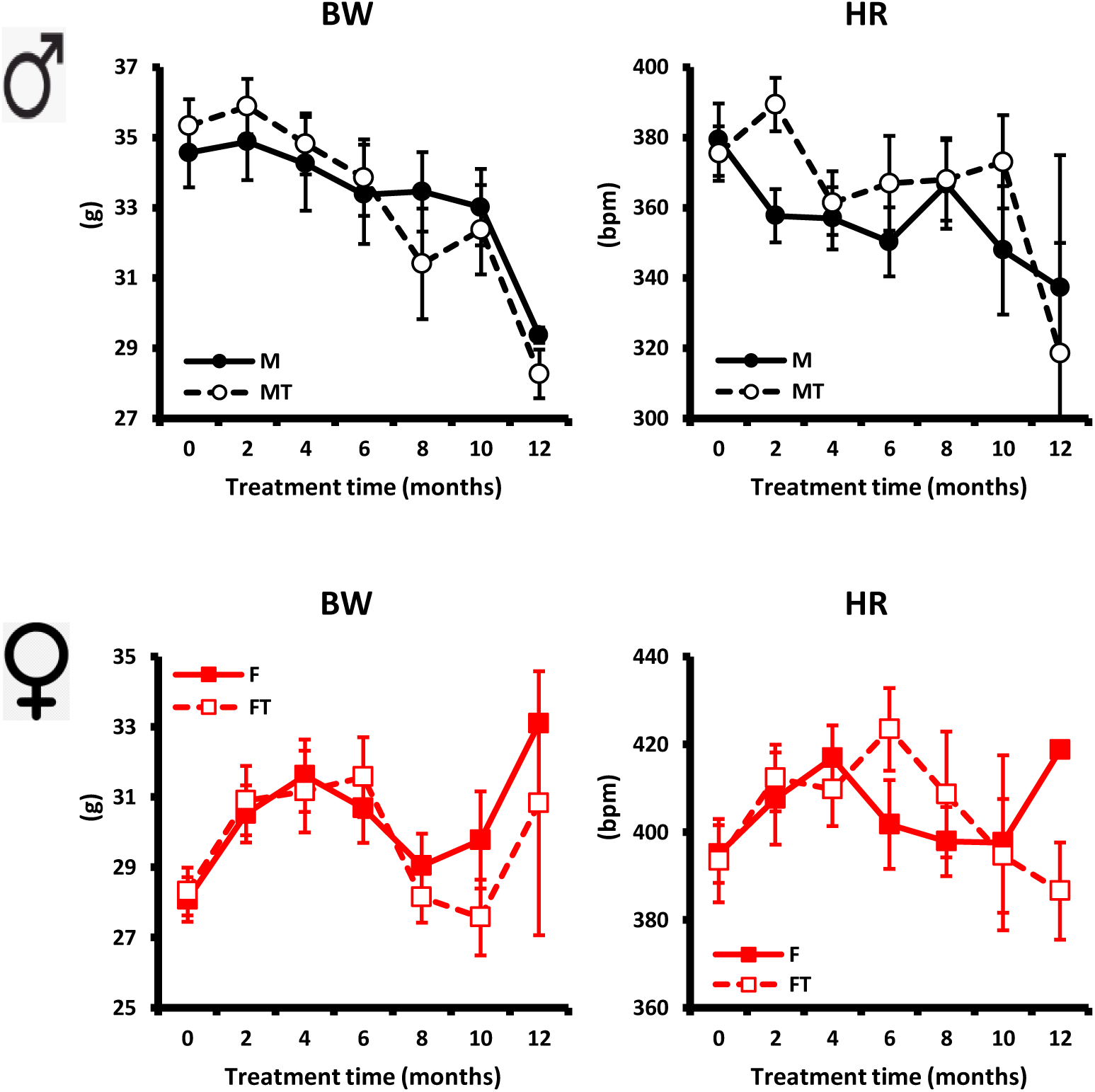
Body weight and heart rate. BW = body weight; HR = heart rate. Mean ± SEM; $ p<0.05 for Group-time interaction; post-hoc test: # (red) p<0.05 F vs. FT; # (black) p<0.05 M vs. MT.

### Survival Rate

During the 12-month of observation period, 67 out of 120 mice were deceased by natural causes, and remaining 53 mice were sacrificed for fresh tissue samples. The actual cause of deaths was not clear, but assumed related to age-associated diseases such as cancer, stroke, heart failure or other organ failures. Autopsy results provided some plausible causes but not definite. Fig.11 plots the Kaplan Meier survival curves of the four groups.

**Figure 11:**
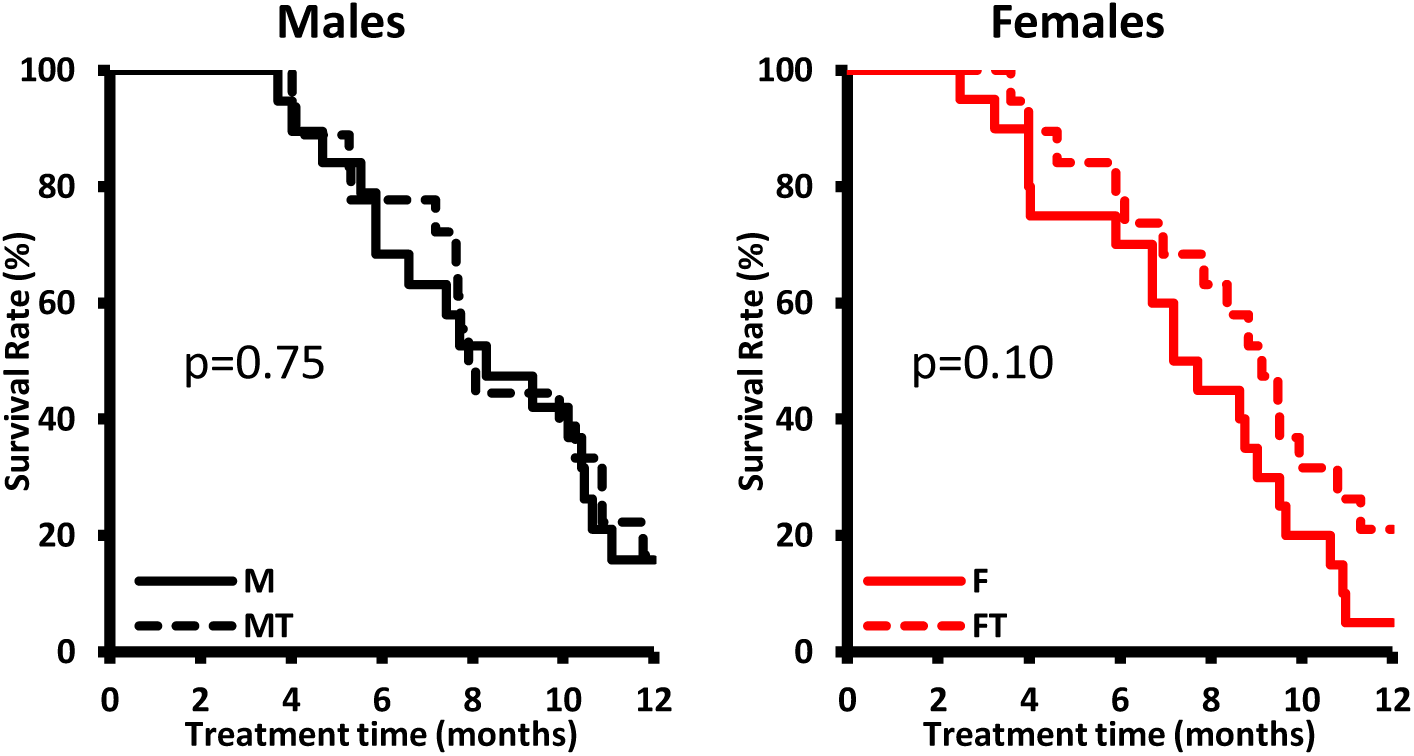
Cumulative survival rate for male (left panel) and female (right panel) mice. It was not significantly different between M and MT groups, and between F and FT groups, respectively.

Accumulated Survival Rate was not significantly different between M and MT groups (p = 0.75) and between F and FT groups (p = 0.10). The median survival time was extended by 0.6 month in MT group compared to M group; and by 1.0 month in FT group compared to F group.

### Autopsy Findings

All 120 mice were underwent autopsy examinations. The prevalence of dermatitis, abdominal tumors, chest tumors and stroke that calculated from autopsy results was shown Fig.12. The prevalence of dermatitis and stroke were significantly lower in MT compared to M group; and FT group compared to F group. However, the prevalence of chest tumors was significantly higher in MT and FT groups compared to M and F groups, respectively. The prevalence of abdominal tumors was similar in MT and FT groups compared to M and F groups, respectively.

**Figure 12:**
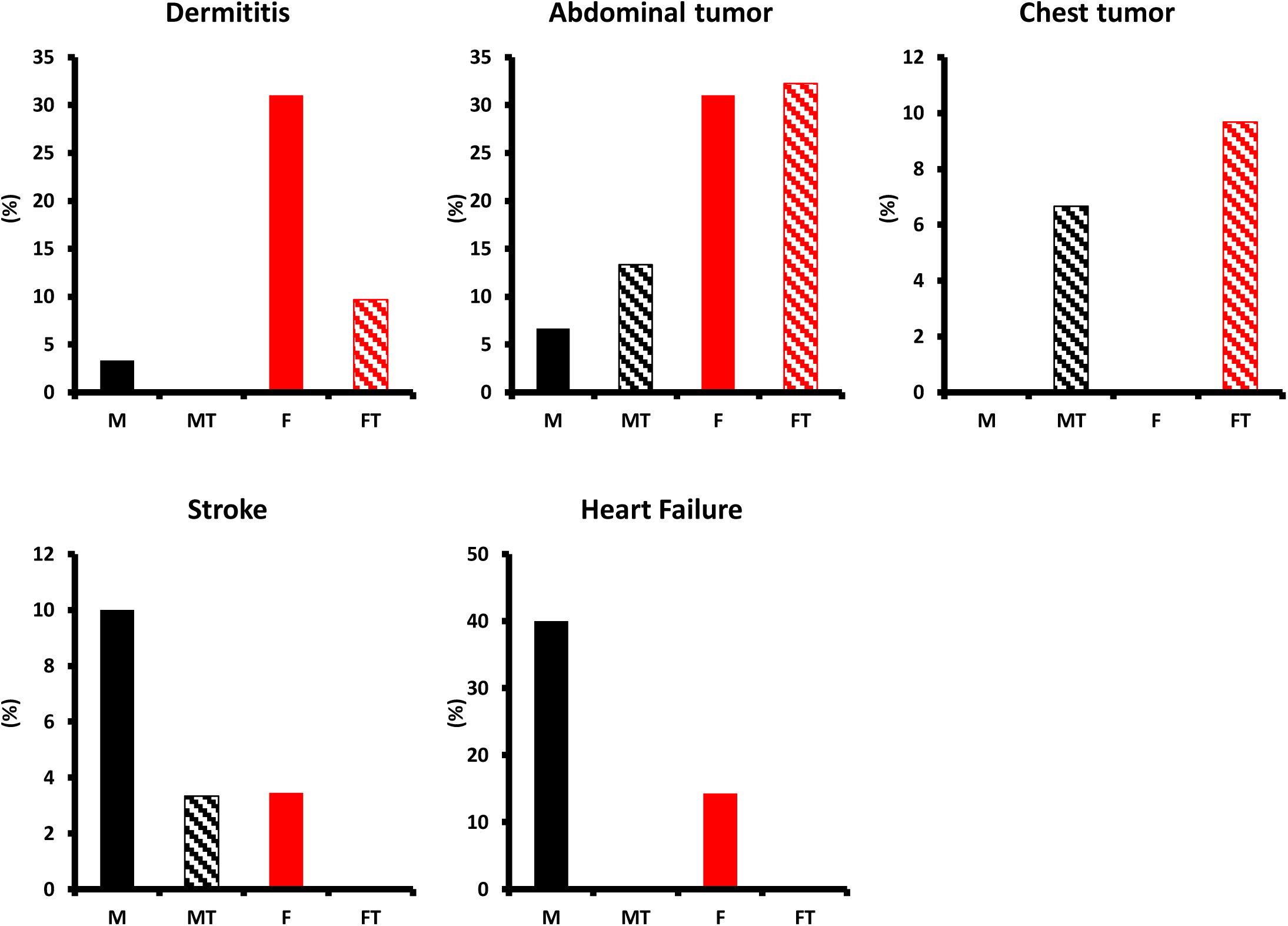
Prevalence of dermatitis, abdominal tumors, chest tumors, stroke and heart failure reported from autopsy. Prevalence was shown as a percent.

## DISCUSSION

In this study, we have shown, for the first time, the beneficial effects of long-term whole body exposure of low dose Near Infrared Light (NIR) treatments on cardiovascular aging, gait and frailty in a normal mouse of both genders. A very brief (2 min/day) exposure to light that was started at middle ages of mice and carried on till very old ages (human equivalent of 54 to 90 years of age) significantly altered the course of age-associated cardiovascular remodeling, improved skeletomuscular function and improved overall health in these mice. This is the first *in vivo* long-term study, to the best of our knowledge that demonstrate the remarkable effects of PBM in the context of aging in both genders. Not only male and female mice showed a very different cardiovascular aging process unexpectedly (as shown in Supp. Table.1), but also they showed a very different response to the same type of PBM exposure.

Recently, we have reported that PBM attenuates cardiovascular remodeling and extends the lifespan in a mouse model of accelerated aging [6]. In that study, 14-month old male AC8 mice and wild type siblings (C57 background) were exposed daily to PBM for the first 3 months and again the last 1 month during an 8-month observation period. A mere 3-month of daily exposure to NIR for briefly (2 min) was sufficient to make a remarkable difference in overall health of sick mice. As a continuity, here we showed again that such exposure to NIR were potent enough to make a difference in overall health of a normal mouse in the absence of a clinical ailment.

Whole body exposure to PBM, not only attenuated the age-associated cardiovascular remodeling, but also attenuated the age-associated deterioration in gait, frailty and thermoregulation, and reduced the prevalence of dermatitis, stroke and heart failure. Although, the increased prevalence of chest tumors gave a cautionary note, overall, we didn’t observe a clear toxicity of PBM in cardiovascular system associated with a-year-long daily exposure.

In our previous findings [6], we observed significant perturbations in TGF-β_1_ levels in serum with PBM treatments. Considering the critical role TGF-β in cellular degeneration, fibrosis, inflammation, regenerative capacity and metabolic functions in humans and mouse alike [34, 35], it is clear that TGF-β_1_ signaling is part of the mechanistic pathways of PBM. However, others signaling pathways such as ATF-4, NFκB, VEGF, and PI3K may also play a role that requires further investigations.

During last decade, PBM has shown promising outcomes in many ailments [36, 37]. PBM is believed to have significant roles in modulating biological processes/pathways to promote cellular viability and restoring the cellular functions following injury through multiple signaling pathways [4]. However, the precise mechanisms for the therapeutic effects of PBM in cardiovascular aging we observed here remain to be elucidated and are outside the scope of current study. In addition, duo to PBM still in its infancy, PBM parameters used in the published literature is varying greatly. The lack of rigorous dosimetry and lack of attention to precise biological targets has plagued the field with inconsistent clinical outcomes. Encouraged by the remarkable outcomes in our previous PBM study [6], we used the same PBM parameters and treatment method in this study. However, further studies on PBM dose-escalation and temporal kinetics of cellular and molecular changes would be needed to formulate a best strategy for this new phenomenon.

During the latter phase of lifespan, mice exhibit all the characteristics of acceleration in cardiovascular aging process. As shown in this study, mice exhibited a significant cardiac chamber dilatation, functional deterioration, hypertrophy and diastolic dysfunction, and stiffening the aortic wall, in addition to deterioration in Gait, frailty and thermoregulation. Increased prevalence of tumors, dermatitis, stroke and heart failure leads to premature deaths. Surprisingly, there are some gender difference in the aging process: in contrast to a steady decline in body mass and heart rate in males, a biphasic fluctuation was observed in females; in contrast to the mainly LV functional remodeling in males, a LV structural remodeling was dominant in females, as a result, while the blood perfusion of vital organs was suffered in males, it was insulated in females; in contrast to high prevalence of stroke and heart failure in males, dermatitis and abdominal tumors were the main feature in females, so on. PBM therapy generated similar effects in some but not all parameters in both genders. While the effects of PBM on heart in males were felt as an improved blood perfusion of whole body, it was largely muted in females even PBM attenuated the LV structural remodeling. A further investigation into the gender difference would be very important in the context of future clinical translation of PBM therapy.

As proof of concept, this study focused mainly on the changes in cardiovascular system by design and at lesser extent on brain-limb coordination, frailty and other organ systems. Effects of PBM on other organ systems, skin and tumor growth can also be further explored. Investigations into precise mechanisms were also limited at this stage. Gender difference in response to PBM therapy added an interesting twist to existing clinical translation strategies.

In conclusion, we showed that PBM can be a simple, cheap and practical therapeutic addition to the treatment of cardiovascular aging, and potentially for also tumors, dermatitis and other ailments if proven further.

## ACKNOWLEDGEMENTS

This research was fully supported by the NIA IRP.

## AUTHOR CONTRIBUTIONS

IA designed the experiments, performed the experiments, supervised the experiments, and wrote the manuscript; SS performed the experiments; KC performed the experiments; CM performed the statistical analysis; PA reviewed the manuscript; EL supervised the experiments. All authors approved the final version of the manuscript.

