## supplement for "The Therapeutic Effects of Long-term Photobiomodulation on Aging in Mice"

Age effect and age x gender interaction for male and female untreated groups were analyzed. Only the parameters that showed different age effects between male and females and parameters that showed a significant age-gender interaction ($p<0.05$) were listed in Table.1. P values in yellow indicate a trend ($0.05<p<0.10$).

**Table 1**: List of Cardiovascular, Gait and Frailty Parameters changed differently by age and sex

| | Age effect | | Age x Gender interaction |
|---|---|---|---|
| | Male | Female | |
| BW | 0.00 | 0.00 | 0.00 |
| LVM | 0.00 | 0.00 | 0.08 |
| EDV | 0.00 | 0.00 | 0.00 |
| PW | 0.00 | | |
| LAD | 0.00 | 0.00 | 0.00 |
| LADc | 0.10 | 0.00 | 0.00 |
| ESV | | 0.00 | |
| HR | 0.01 | 0.04 | 0.06 |
| SV | 0.00 | 0.00 | 0.00 |
| CO | 0.00 | 0.00 | 0.00 |
| CI | 0.00 | 0.00 | 0.00 |
| E | | 0.00 | 0.02 |
| e' | | 0.00 | 0.00 |
| a' | | 0.09 | |
| e'/a' | 0.02 | | |
| E/e' | 0.05 | 0.03 | 0.07 |
| CAFlow/CO | 0.00 | 0.01 | 0.10 |
| Gait Speed | 0.00 | 0.00 | 0.03 |
| Animal Length | 0.00 | 0.00 | 0.01 |
| Stance time | 0.00 | 0.00 | 0.02 |
| Propel time | 0.00 | 0.00 | 0.06 |
| Stride time | 0.00 | 0.00 | 0.02 |
| Brake time | 0.00 | 0.00 | 0.01 |
| PFI | 0.00 | | 0.08 |
| TFI | 0.04 | | |
| SFI | 0.08 | | |
| Body Temp | 0.03 | | 0.02 |

*Data as p values for age effect for male and females untreated and gender x age interaction.*

**Figure 1:** Cardiac diastolic parameters. E/A = Mitral flow E to A ratio; e'/a' = Mitral tissue Doppler e' to a' ratio; Mean ± SEM; $ $p<0.05$ for Group-time interaction; post-hoc test: # (red) $p<0.05$ F vs. FT; # (black) $p<0.05$ M vs. MT.

**Figure 2-5:** Gait Parameters. Mean ± SEM; $ $p<0.05$ for Group-time interaction; post-hoc test: # (red) $p<0.05$ F vs. FT; # (black) $p<0.05$ M vs. MT.

**Figure 6:** Aortic pulse wave velocity (PWV). Mean ± SD. There are statistically significant age, treatment and age*treatment effects between PBM treatment and no-treatment groups.

### Fig.1

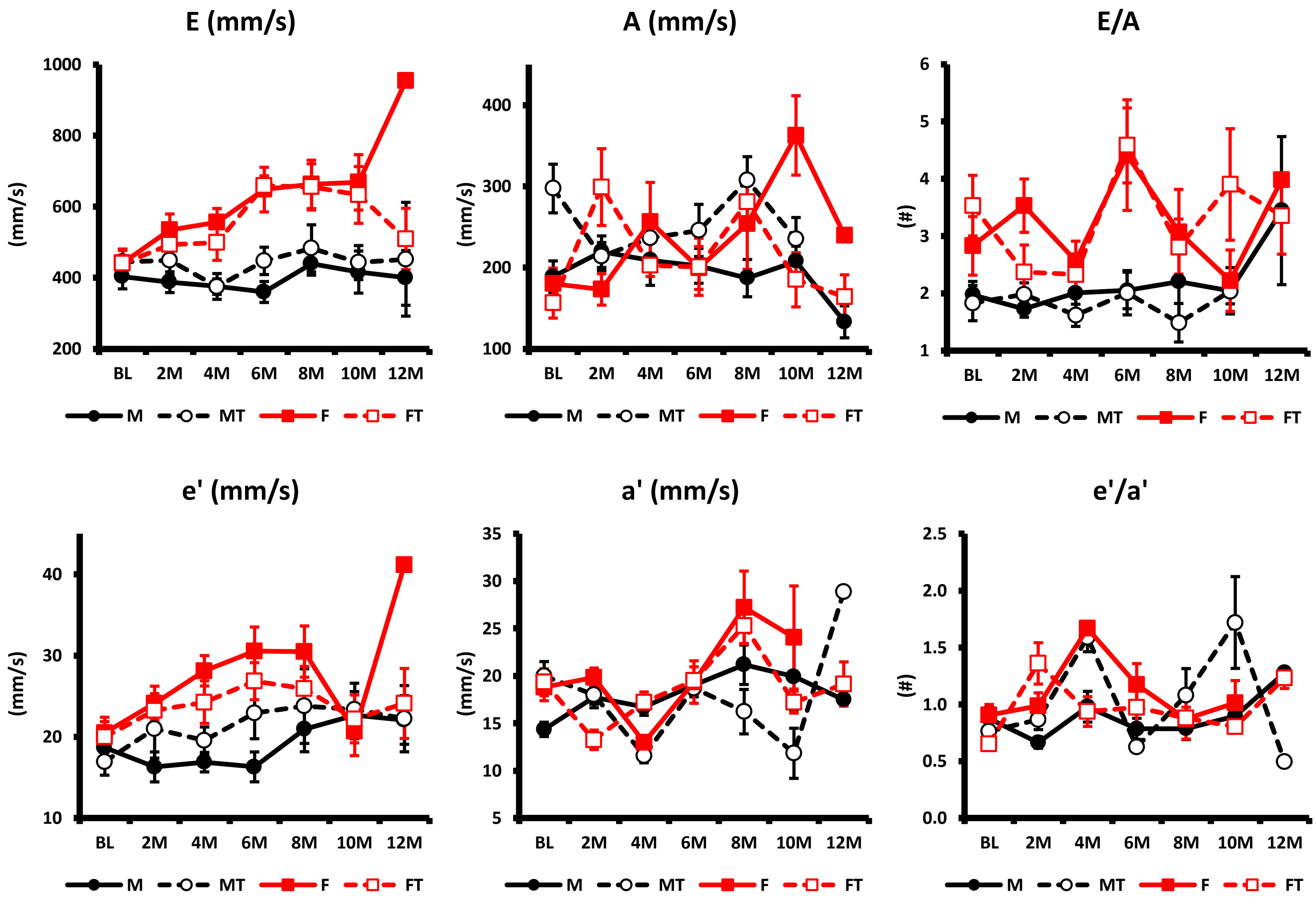


**Figure 1:** Cardiac diastolic parameters. E/A = Mitral flow E to A ratio; e'/a' = Mitral tissue Doppler e' to a' ratio; Mean ± SEM; # (red) p<0.05 F vs. FT; # (black) p<0.05 M vs. MT.

Fig.2A

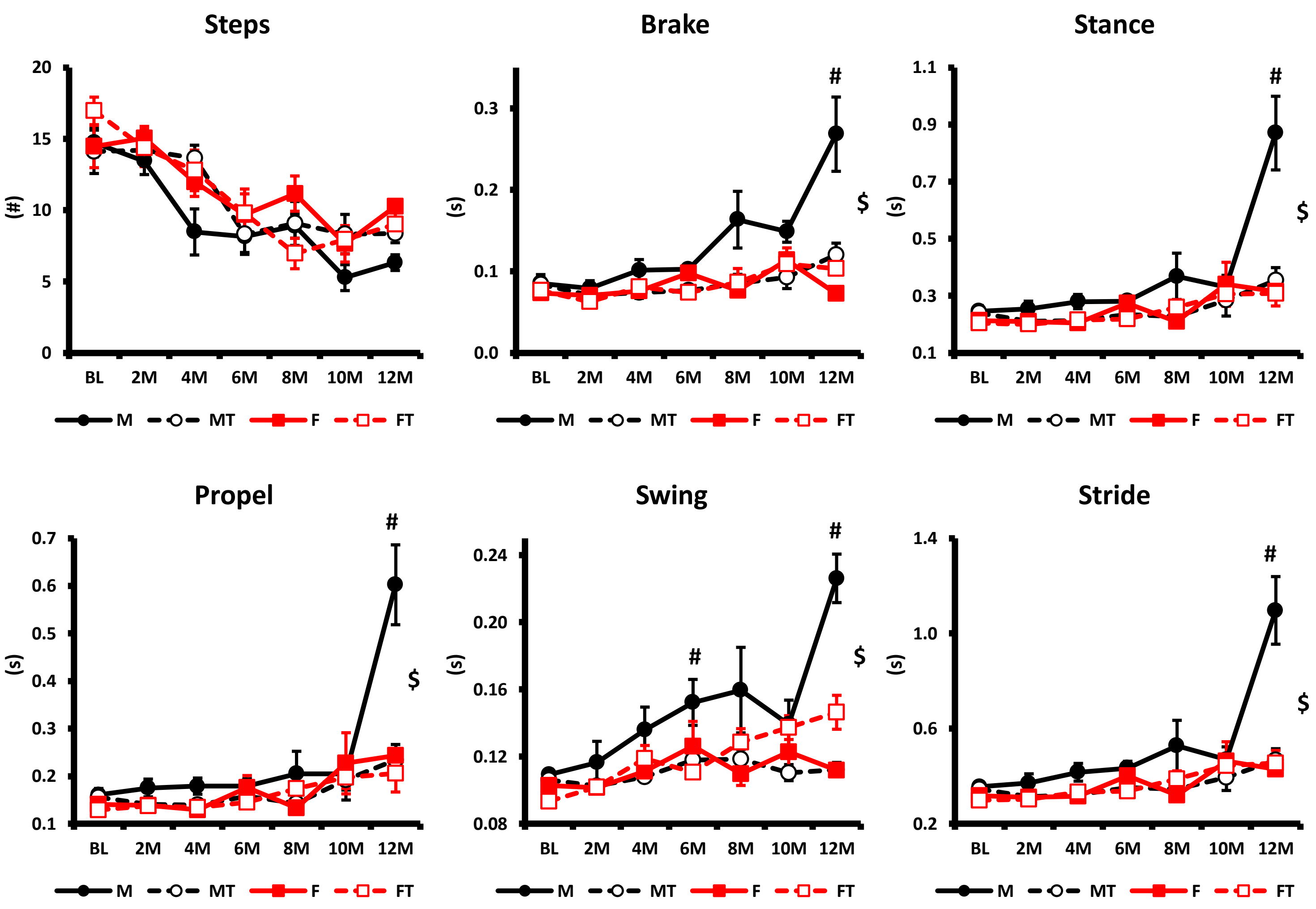


**Figure 2:** Gait Temporal Parameters. Mean ± SEM; # (red) p<0.05 F vs. FT; # (black) p<0.05 M vs. MT.

Fig.2B

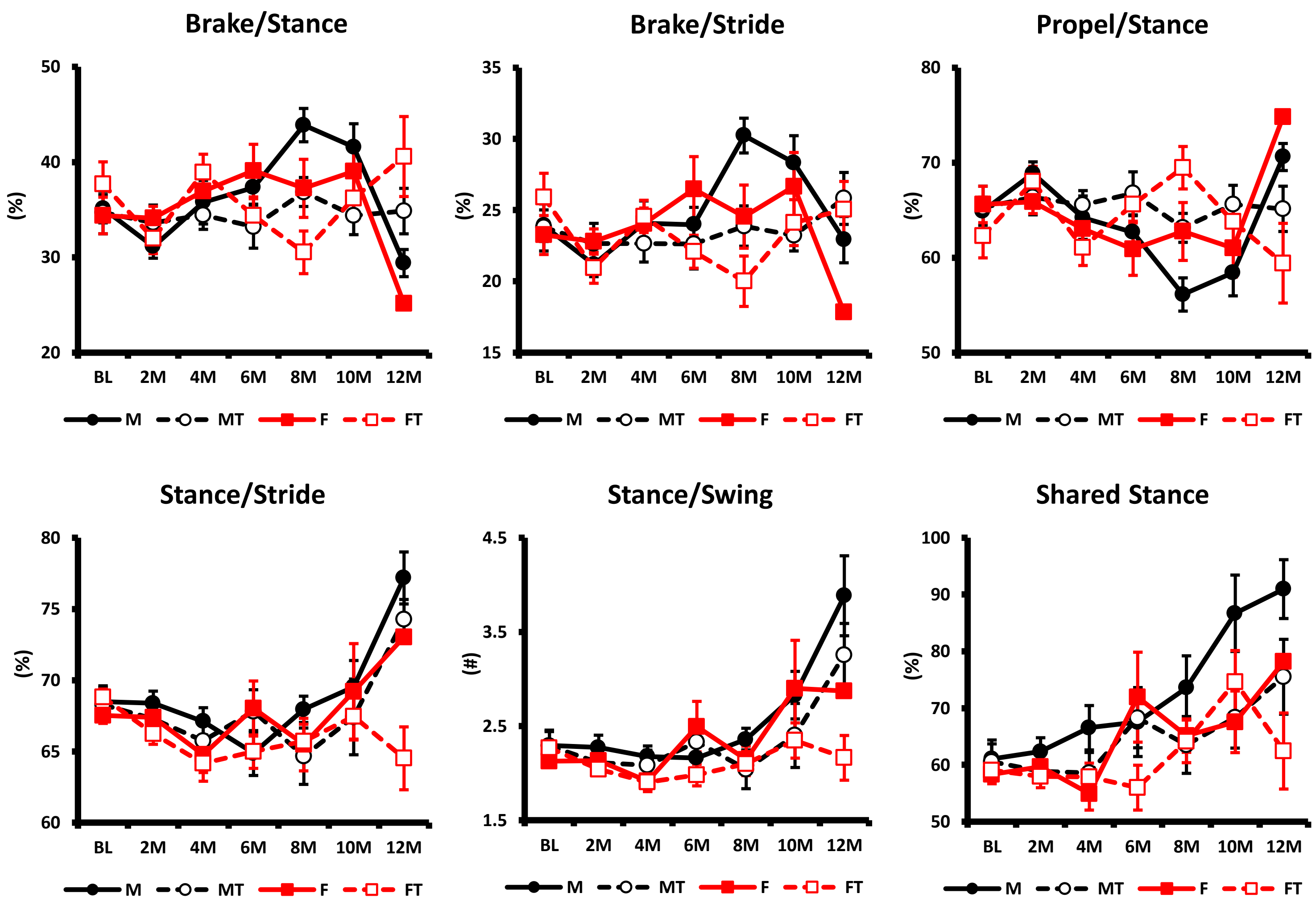


**Figure 2:** Gait Temporal Parameters. Mean ± SEM; # (red) p<0.05 F vs. FT; # (black) p<0.05 M vs. MT.

Fig.2C

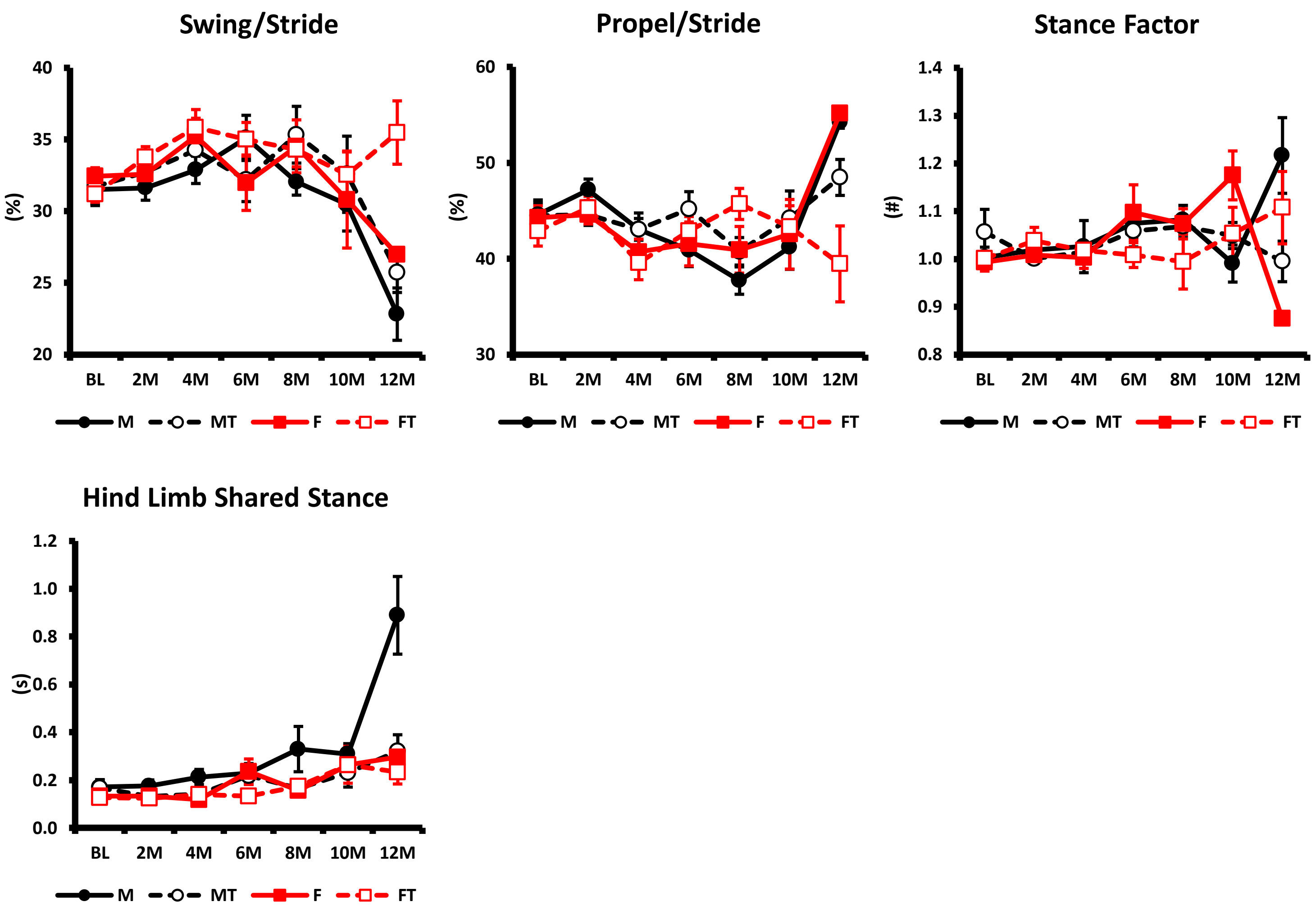


**Figure 2:** Gait Temporal Parameters. Mean ± SEM; # (red) p<0.05 F vs. FT; # (black) p<0.05 M vs. MT.

Fig.3A

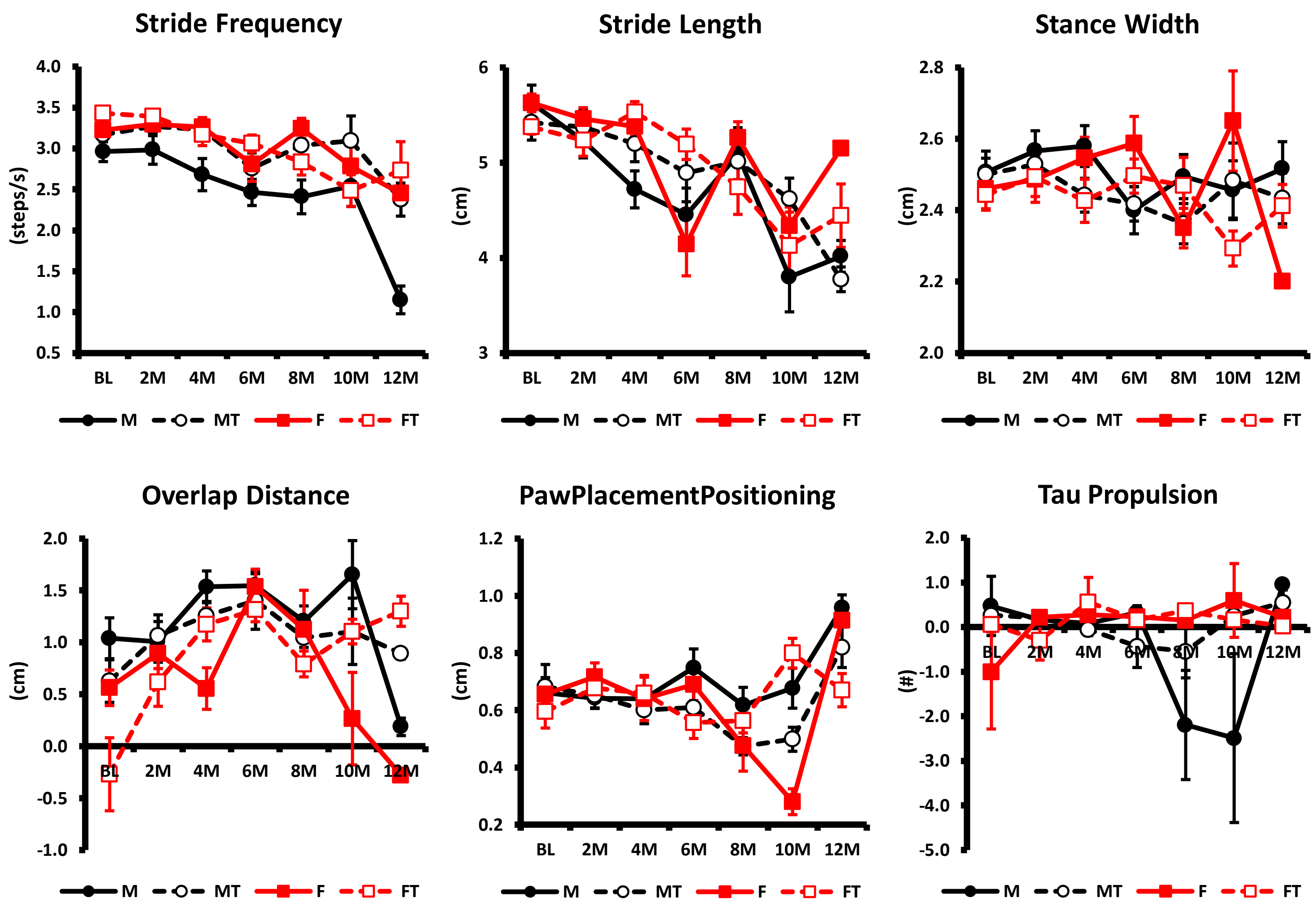


**Figure 3:** Gait Spatial Parameters. Mean ± SEM; # (red) p<0.05 F vs. FT; # (black) p<0.05 M vs. MT.

### Fig.3B

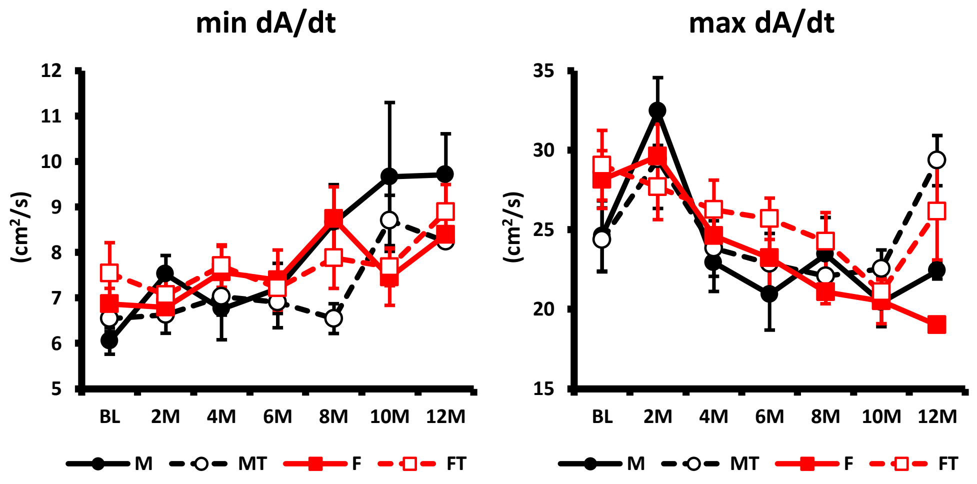


**Figure 3:** Gait Spatial Parameters. Mean ± SEM; # (red) $p<0.05$ F vs. FT; # (black) $p<0.05$ M vs. MT.

### Fig.4A

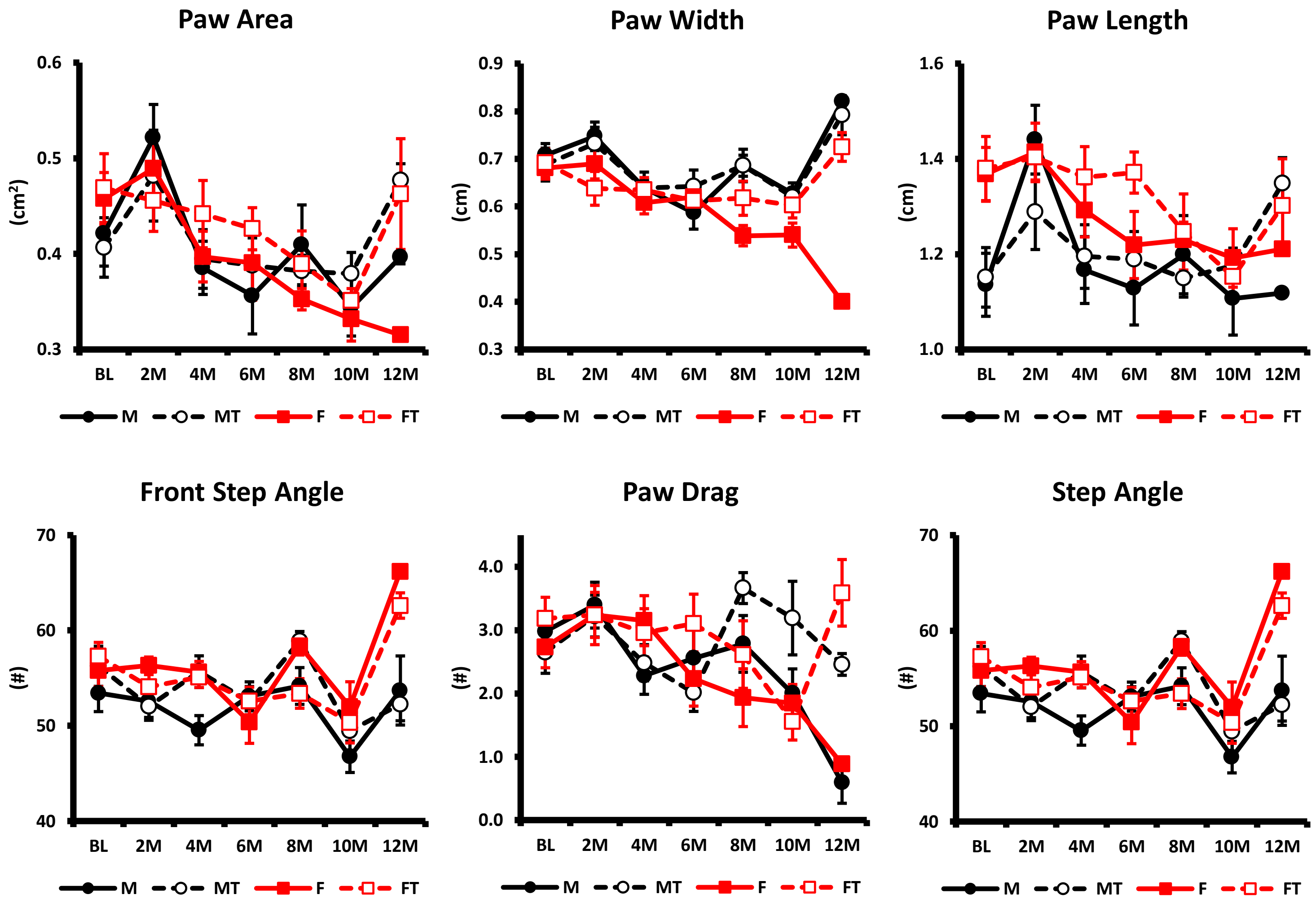


**Figure 4:** Gait Postural Parameters. Mean ± SEM; # (red) p<0.05 F vs. FT; # (black) p<0.05 M vs. MT.

Fig.4B

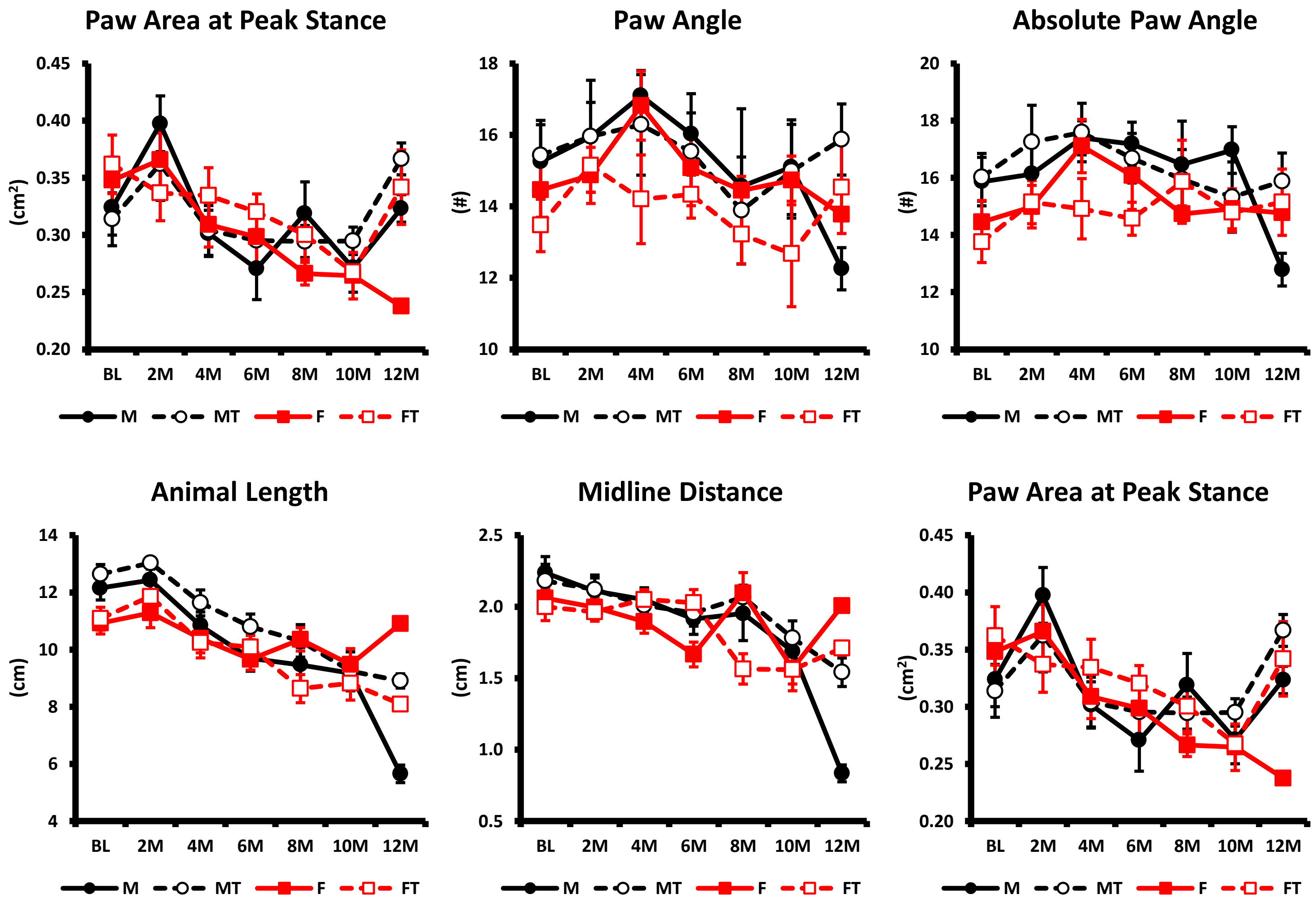


**Figure 4:** Gait Postural Parameters. Mean ± SEM; # (red) $p<0.05$ F vs. FT; # (black) $p<0.05$ M vs. MT.

### Fig.4C

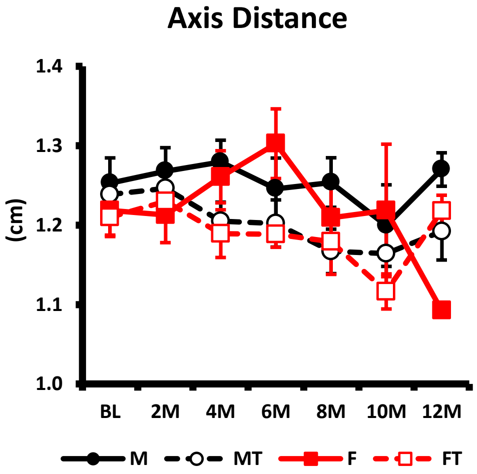


**Figure 4:** Gait Postural Parameters. Mean ± SEM; # (red) p<0.05 F vs. FT; # (black) p<0.05 M vs. MT.

### Fig.5A

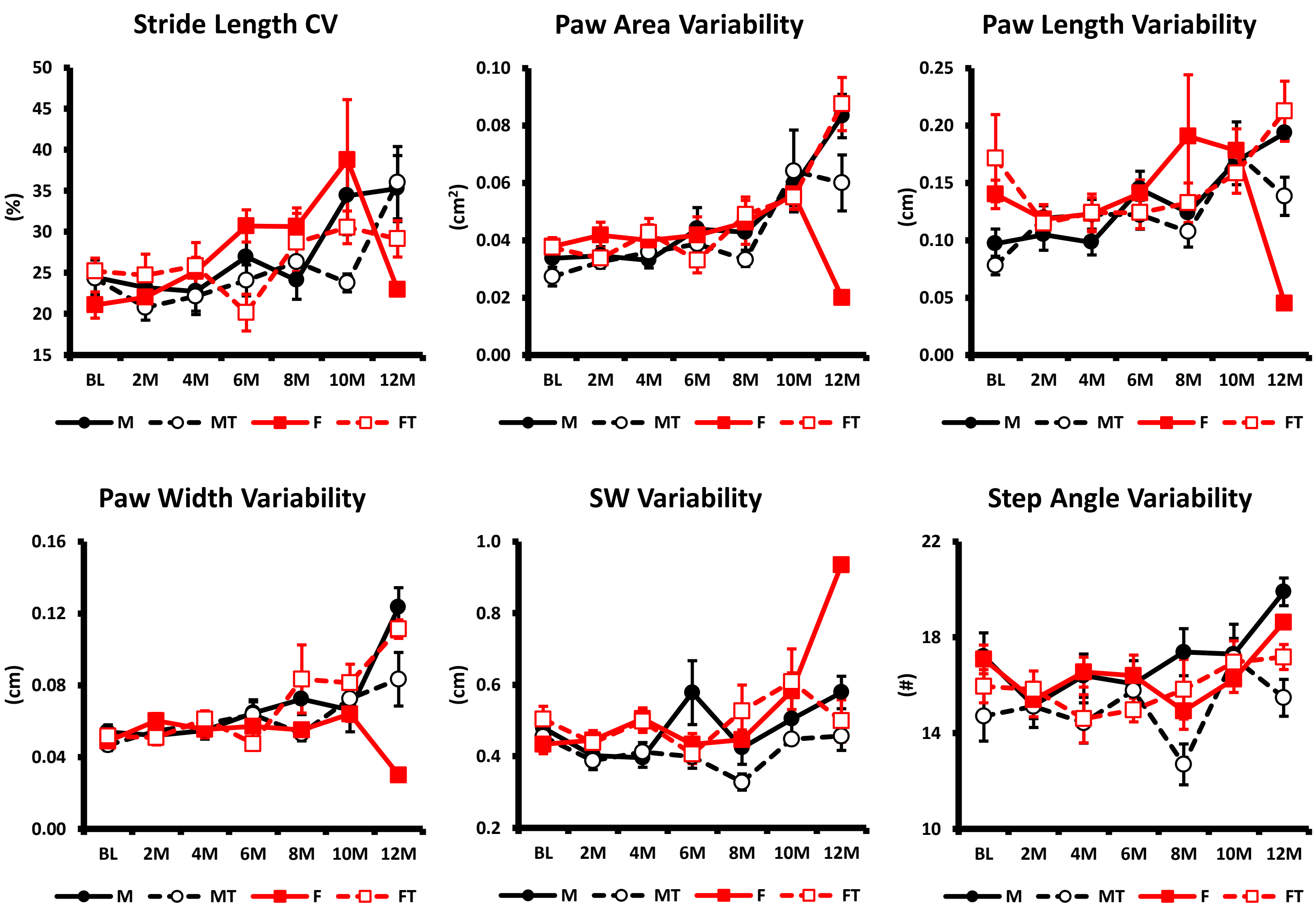


**Figure 5:** Gait Intra-individual Variability Parameters. Mean ± SEM; # (red) p<0.05 F vs. FT; # (black) p<0.05 M vs. MT.

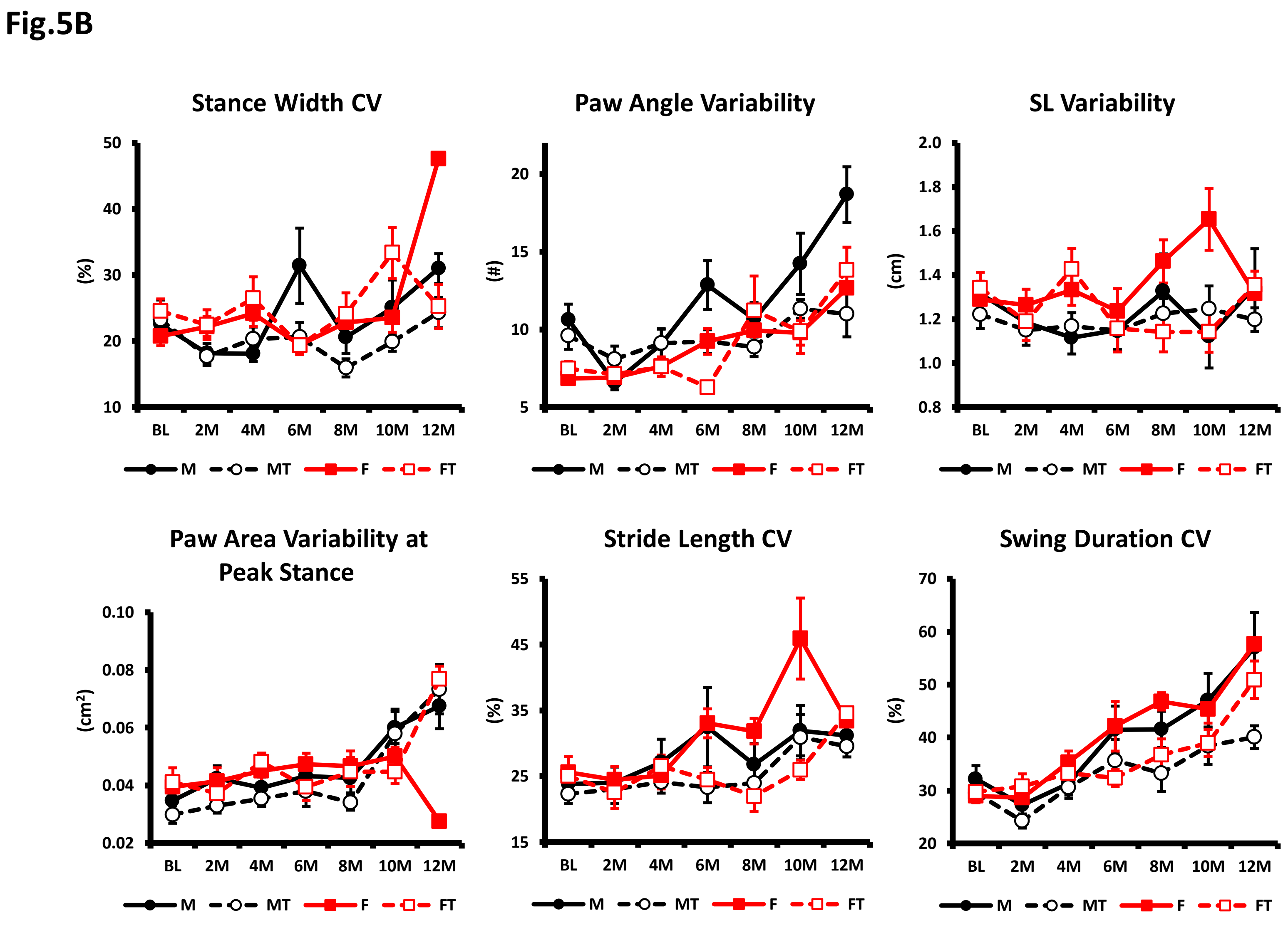


**Figure 5:** Gait Intra-individual Variability Parameters. Mean ± SEM; # (red) $p<0.05$ F vs. FT; # (black) $p<0.05$ M vs. MT.

**Fig.5C**

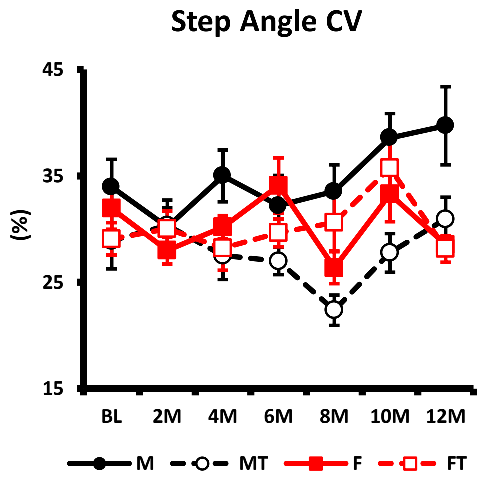


**Figure 5:** Gait Intra-individual Variability Parameters. Mean ± SEM; # (red) $p<0.05$ F vs. FT; # (black) $p<0.05$ M vs. MT.

**Fig.6**

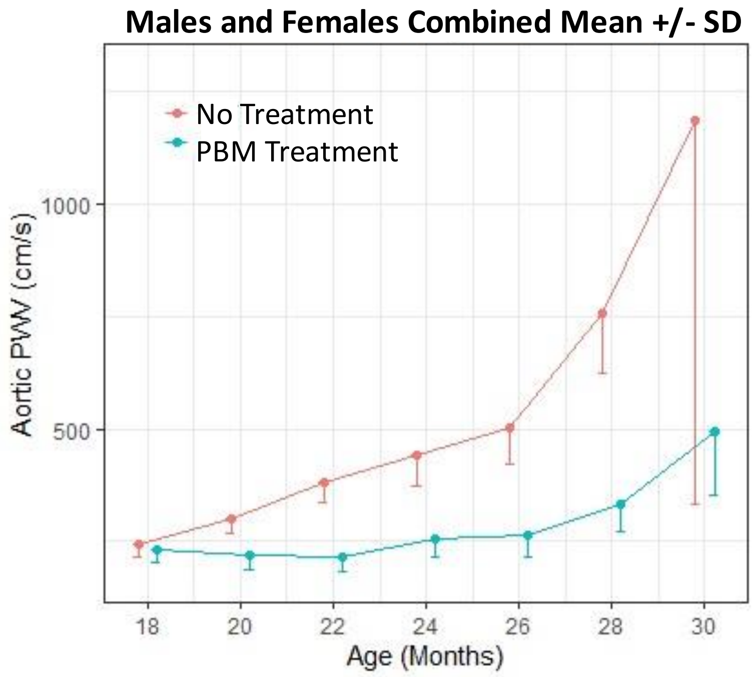


**Figure 6:** Aortic pulse wave velocity (PWV). Mean± SD. There are statistically significant age, treatment and age*treatment effects between PBM treatment and no-treatment groups.
